# Adropin confers neuroprotection and promotes functional recovery from ischemic stroke

**DOI:** 10.1101/2021.09.16.460662

**Authors:** Changjun Yang, Bianca P. Lavayen, Lei Liu, Brian D. Sanz, Kelly M. DeMars, Jonathan Larochelle, Marjory Pompilus, Marcelo Febo, Yu-Yo Sun, Yi-Min Kuo, Mansour Mohamadzadeh, Susan A. Farr, Chia-Yi Kuan, Andrew A. Butler, Eduardo Candelario-Jalil

## Abstract

Adropin is a highly-conserved peptide that has been shown to preserve endothelial barrier function. Blood-brain barrier (BBB) disruption is a key pathological event in cerebral ischemia. However, the effects of adropin on ischemic stroke outcomes remain unexplored. Hypothesizing that adropin exerts neuroprotective effects by maintaining BBB integrity, we investigated the role of adropin in stroke pathology utilizing loss- and gain-of-function genetic approaches combined with pharmacological treatment with synthetic adropin peptide. Stroke decreased endogenous adropin levels in the brain and plasma. Adropin treatment or transgenic adropin overexpression robustly reduced brain injury and improved long-term sensorimotor and cognitive function in young and aged mice subjected to ischemic stroke. In contrast, genetic deletion of adropin exacerbated ischemic brain injury. Mechanistically, adropin neuroprotection depends on endothelial nitric oxide synthase and is associated with reduced BBB permeability and neuroinflammation. We identify adropin as a novel neuroprotective peptide with the potential to improve stroke outcomes.

## Introduction

Ischemic stroke is a common cause of death and long-term disability in adults that is expected to increase in prevalence due to an aging population (*1*). The only FDA-approved pharmacological treatment for acute stroke, recombinant tissue plasminogen activator (tPA), is provided to a small percentage of stroke patients due to a narrow therapeutic time window (*2*). Results from animal and human studies indicate that the passage of tPA into the ischemic brain due to damage to the blood-brain barrier (BBB) after stroke can result in direct neurotoxicity (*3*). This can increase the incidence of intracerebral hemorrhage (*4*), further limiting its therapeutic use. Developing novel therapeutic strategies against neurovascular injury that lack limiting side effects is thus an imperative for treating ischemic stroke.

Studies from our group and others demonstrated that adropin exerts beneficial effects in preserving endothelial permeability *in vitro* and *in vivo* (*5-7*). Adropin is encoded by the *energy homeostasis associated gene* (*Enho*) (*8*) and is a potential regulator of endothelial function (*5*). The polypeptide is composed of 76 amino acids and is described as consisting of a secretory signal peptide (adropin^1-33^) and a putative secreted domain (adropin^34-76^) (*8*). However, the mechanisms of peptide secretion are poorly understood. There is evidence that adropin is a membrane-bound protein and is highly expressed in the brain (*8, 9*). Adropin is expressed in vascular endothelial cells and appears to play an essential role in regulating endothelial function (*5, 6, 10*). Adropin stimulates angiogenesis, blood flow, and capillary density in a mouse model of hind limb ischemia (*5*). Treatment with adropin also reduced endothelial permeability, increasing vascular endothelial growth factor receptor 2 (VEGFR2) and its two downstream effectors phosphatidylinositol-3-kinase (PI3K)/Akt- and extracellular signal-regulated kinases 1/2 (ERK1/2)-dependent eNOS <u>signaling pathways (</u>*<u>5</u>*).

Our previous findings showed that adropin protects brain endothelial cells exposed to hypoxia and low glucose *in vitro* (*6*). Expression of *Enho* was dramatically downregulated in rat brain microvascular endothelial cells exposed to ischemia-like conditions (*6*). On the other hand, adropin treatment significantly attenuated the increased permeability of the endothelial monolayer in a concentration-dependent manner (*6*). In line with these findings, Yu *et al*. found that adropin treatment could preserve BBB integrity in mice subjected to intracerebral hemorrhage through the Notch1/Hes1 pathway (*7*). There is also emerging evidence of low levels of serum adropin correlating with endothelial dysfunction and severity of coronary heart disease (*11, 12*). Adropin protein levels in the brain and plasma decrease with the aging of rats (*13*), mice (*14*), and humans (*15, 16*). A recent study found lower serum adropin levels in patients with acute ischemic stroke compared to controls (*17*).

Considering that aging-associated endothelial dysfunction and neurovascular damage play a crucial role in ischemic stroke pathology and that adropin reduces endothelial permeability (*5-7*), we hypothesized that adropin plays a protective role in the context of ischemic stroke by maintaining BBB integrity. This possibility has not been explored in preclinical models of stroke. Here we investigated the effects of adropin on anatomical and functional outcomes following experimental ischemic stroke in mice. Utilizing complementary pharmacological and genetic approaches, we identified novel molecular mechanisms of protection by adropin against ischemic brain injury. We found that post-ischemic treatment with synthetic adropin peptide or transgenic overexpression of endogenous adropin produces long-lasting neuroprotection in young and aged animals. Transgenic overexpression of adropin provides robust neuroprotection, while genetic deletion of adropin in *Enho*^*-/-*^ mice significantly exacerbates ischemic brain injury in both male and female mice. Adropin treatment significantly reduces stroke-induced loss of tight junction proteins and BBB damage. The protective effects of adropin in cerebral ischemia depend on eNOS activation, which sheds new light on the actions of adropin in the context of ischemic brain damage. The promising neuroprotective effects of adropin reported here would set the stage for expanded preclinical work on the efficacy of this newly discovered endogenous peptide in cerebral ischemia and other types of brain injury associated with BBB damage. Adropin therapy could emerge as a novel and much-needed approach to reduce the devastating consequences of neurovascular injury after stroke.

## Results

### Adropin is expressed in endothelial cells and neurons in the mouse brain and highly vascularized peripheral organs

Double immunofluorescence staining analysis showed that adropin is abundantly expressed in endothelial cells and neurons in the mouse cerebral cortex (**Figure S1A and S1B**). Minimal adropin immunoreactivity was detected in GFAP or Iba1 immunoreactive cells (data not shown), in agreement with our recent finding that adropin is expressed in neuronal and endothelial cells using the adropin-internal ribosomal entry sequence (IRES)-Cre-tdTomato reporter mouse (*14*). In addition to the cell types expressing adropin in the brain, we also investigated the expression of adropin in different organs of adult male C57BL/6J naïve mice. As shown in **Figure S1C through S1E**, immunoblotting image, densitometric, and heatmap data indicated that adropin is highly expressed in blood vessel-enriched organs, including brain, liver, kidney, and lung, moderately expressed in spleen and intestines, and poorly expressed in the heart and skeletal muscle.

### Ischemic stroke induces loss of endogenous adropin levels in the brain and plasma

To determine whether stroke alters adropin levels in the brain and plasma in adult male mice subjected to pMCAO, we quantified adropin protein levels in the brain by western blotting. We measured plasma adropin concentrations using a commercially available ELISA kit. We found that stroke induced a significant loss of endogenous adropin levels in the ipsilateral (ischemic) cerebral cortex in a time-dependent manner (**Figure 1A and 1B**). Also, a decrease of adropin in plasma was observed at 48h in mice following pMCAO (**Figure 1C**). These findings suggest that maintenance of brain and/or plasma adropin levels might be helpful to minimize stroke injury.

**Figure 1.**
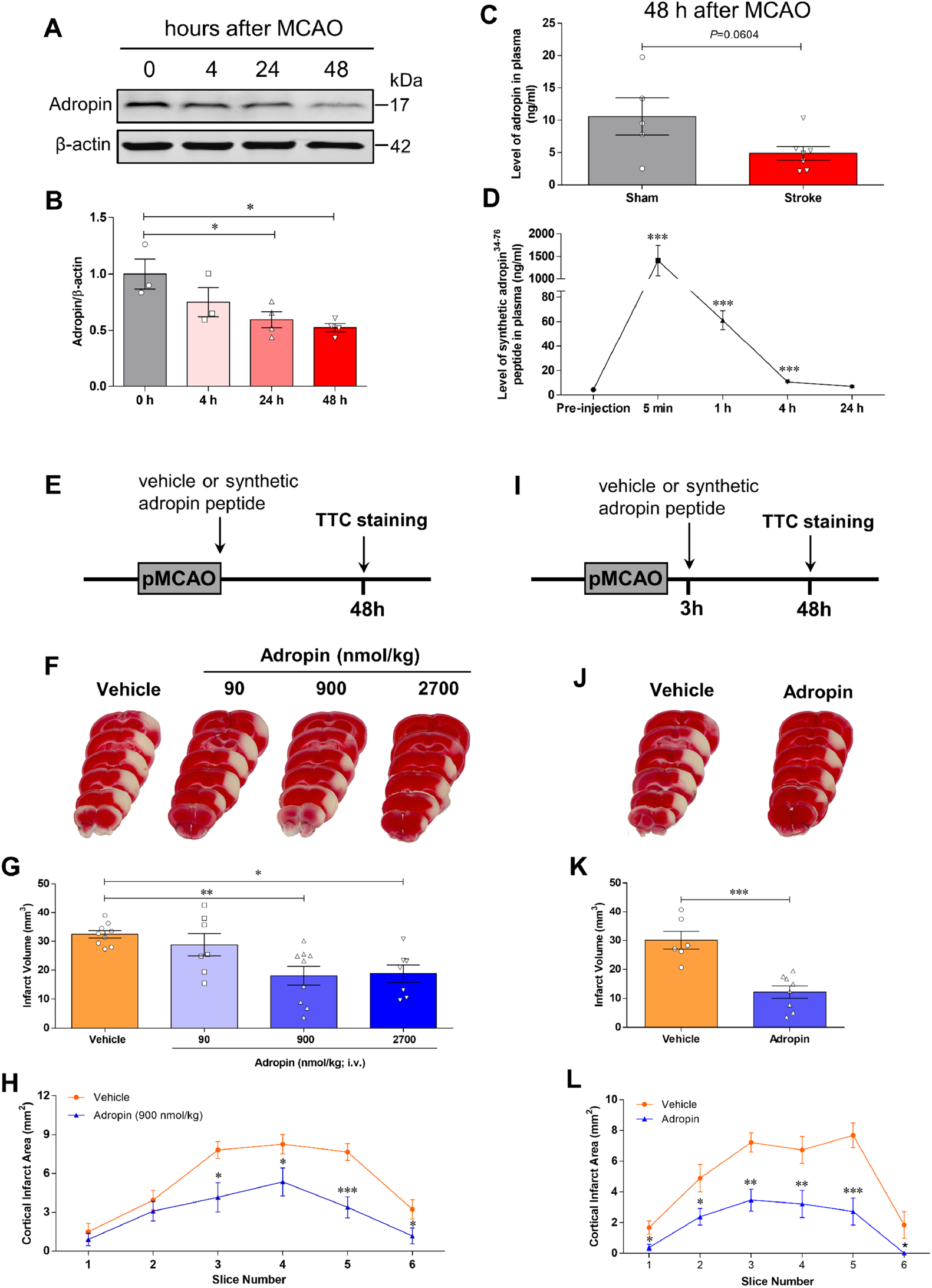
Stroke decreases endogenous adropin levels in the brain and plasma, and post-ischemic treatment with synthetic adropin peptide reduces infarct volume in mice subjected to pMCAO. **A**, Western blot shows stroke-induced loss of endogenous adropin levels in the ipsilateral cerebral cortex in a time-dependent manner. **B**, Densitometric analysis shows that brain adropin levels are significantly reduced from 24 to 48 hours after left middle cerebral artery occlusion. One-way ANOVA with Bonferroni post-tests, \**P*<0.05, n=3-4 per group. **C**, Plasma adropin level was reduced after stroke as assessed by a commercially available adropin ELISA kit. Unpaired t-test, *P*=0.0604, n=5-7 per group. **D**, Dynamic changes of synthetic adropin peptide in plasma. Plasma was collected before (pre-injection) and at 5 min, 1 h, 4h, and 24h after one bolus injection with synthetic adropin^34-76^ peptide (900 nmol/kg; i.v.). Levels of synthetic adropin peptide in plasma dropped to almost pre-injection values within 4 hours. One-way ANOVA with Bonferroni post-tests, \*\*\**P*<0.001, n=4 per group. **E**, Treatment timeline for adult mice subjected to permanent middle cerebral artery occlusion (MCAO) with different doses of synthetic adropin peptide. **F**, Representative 2,3,5-triphenyltetrazolium chloride (TTC)-stained coronal sections from vehicle-, 90-, 900- and 2700-nmol/kg adropin-treated mice. **G**, Post-ischemic treatment with synthetic adropin^34-76^ peptide (90-2700 nmol/kg; i.v.) at the onset of pMCAO significantly reduces brain infarct volume in a dose-dependent manner. One-way ANOVA with Bonferroni post-tests, \**P*<0.05, \*\**P*<0.01, n=7-9 per group. **H**, Analysis of cortical infarct area per-1 mm slice shows that the decrease in cortical infarction by 900 nmol/kg adropin is seen throughout the cortex. Two-way ANOVA with Bonferroni post-tests, \**P*<0.05, \*\*\**P*<0.001. Vehicle (n=9), Adropin (n=9). **I**, Timeline for adult mice subjected to pMCAO with 3h delayed synthetic adropin treatment. **J**, Representative TTC-stained coronal sections of vehicle- and adropin-treated mice. **K**, Post-ischemic treatment with synthetic adropin peptide (900 nmol/kg; i.v.) at 3h after pMCAO significantly reduces brain infarct volume. Unpaired t-test, \*\*\**P*<0.001, n=6-8 per group. **L**, Analysis of cortical infarct area per-1 mm slice shows a reduction in cortical infarction by adropin treatment at different levels of the cerebral cortex. Two-way ANOVA with Bonferroni post-tests, \**P*<0.05, \*\**P*<0.01, \*\*\**P*<0.001. Vehicle (n=6), Adropin (n=8).

### Post-ischemic treatment with synthetic adropin peptide reduces acute ischemic brain injury in adult mice following stroke

Next, we assessed the effects of synthetic adropin (adropin^34-76^) on brain infarct size in adult male mice subjected to pMCAO. To determine the elimination of adropin peptide from the circulation, levels were measured by ELISA in mice administered synthetic adropin peptide (900 nmol/kg; i.v.) in plasma collected before (pre-injection) and at 5 min, 1 h, 4 h, and 24 h after injection. The integrity of the synthetic adropin peptide before injection was confirmed by HPLC-MS/MS and western blot (**Figure S2A through S2D**). Levels of intact adropin^34-76^ in plasma had declined to almost pre-injection values within 4 hours **Figure 1D**, indicating a relatively rapid clearance of the synthetic adropin peptide in mice.

Subsequently, we treated mice with different doses of adropin^34-76^ (90-2700 nmol/kg; i.v.) at the onset of pMCAO and brain infarct volume measured at 48h after stroke by TTC staining (**Figure 1E**). Post-ischemic treatment with adropin at the start of cerebral ischemia significantly reduced cortical infarct volume in a dose-dependent manner (**Figure 1F through 1H**). To explore the potential translational value of adropin treatment in ischemic stroke when given after the onset of stroke, we delayed treatment for 3h after the start of pMCAO (**Figure 1I**). Post-ischemic treatment with synthetic adropin peptide using the optimal dose (900 nmol/kg; i.v.) significantly reduced infarction throughout the cerebral cortex (**Figure 1J through 1L**).

### Post-ischemic treatment with synthetic adropin peptide provides long-lasting neuroprotection and improvement of neurological function in adult mice following stroke

To address whether adropin treatment provides anatomical protection and impacts long-term neurological recovery, we intravenously injected adult male mice with either vehicle (0.1% BSA in saline) or synthetic adropin peptide (900 nmol/kg) at the onset of cerebral ischemia. A battery of neurobehavioral tests was performed on vehicle- and adropin-treated mice before (baseline) and at defined time points (24h, 48h, 7d, 14d, and 21d) after pMCAO (**Figure. 2A**). Brain infarct volume was assessed at 21d after stroke by Cresyl violet staining. Post-ischemic treatment with synthetic adropin peptide significantly reduced brain cortical infarction compared to vehicle-treated animals (**Figure 2B and 2C**). Compared to the vehicle group, adropin treatment significantly reduced the time for mice to sense and remove the sticker on the contralateral paw as assessed by the adhesive removal test at early time points (up to 48 h) after pMCAO (**Figure 2D and 2E**). Similarly, sensorimotor deficits and asymmetry impairment were significantly alleviated up to 21d after stroke in the adropin-treated mice compared to the vehicle group, as assessed using the cylinder test, open field, and corner test (**Figure 2F through 2L**). Although stroke resulted in body weight loss, there was no significant difference in the body weight loss between adropin- and vehicle-treated mice over 21 days after pMCAO (**Figure 2M**). Collectively, these data strongly suggest that synthetic adropin peptide has remarkable effects of reducing infarct volume and improving long-term neurological deficits in ischemic stroke.

**Figure 2.**
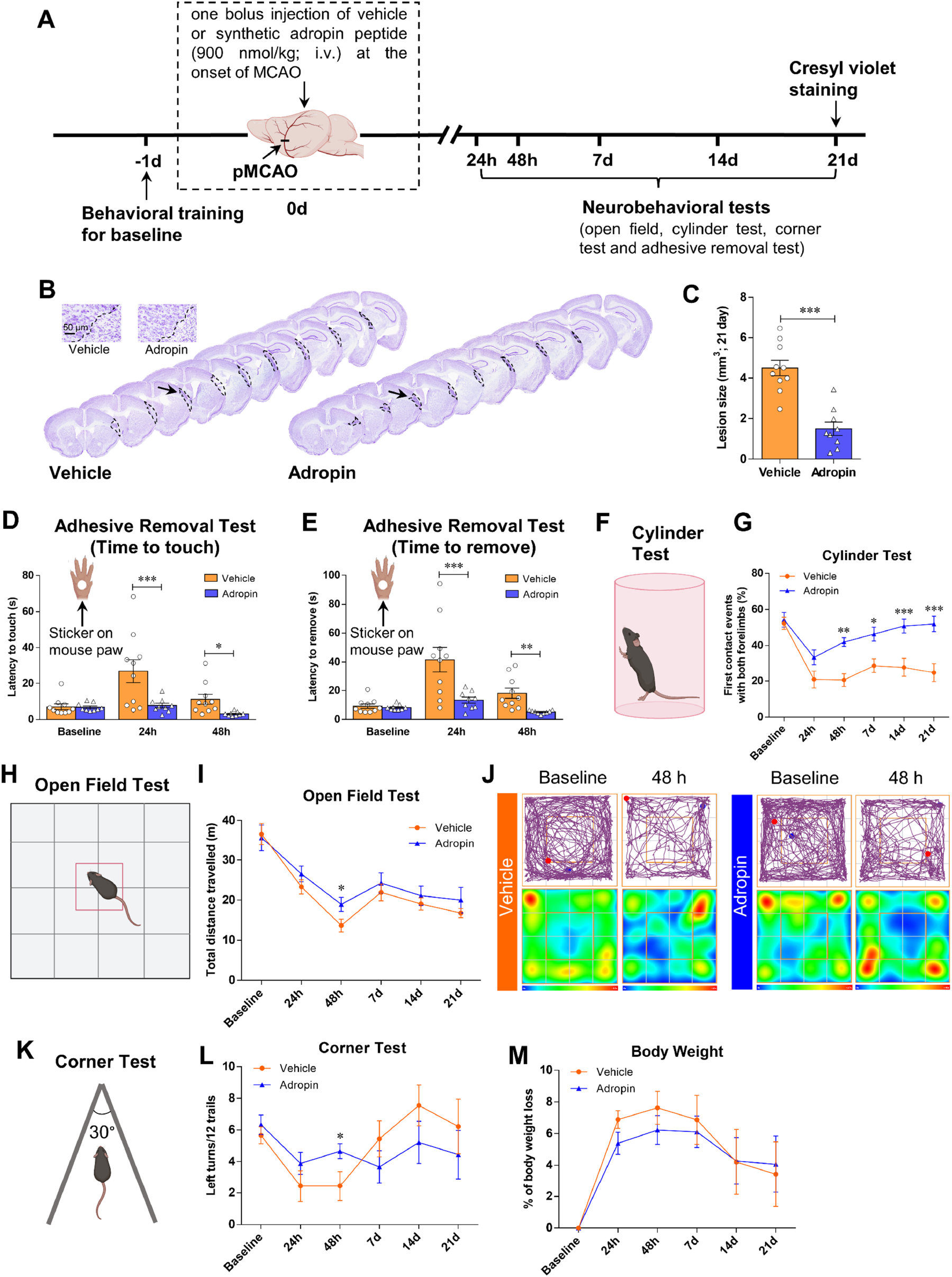
Post-ischemic treatment with synthetic adropin peptide reduces infarct volume and improves long-term neurological deficits in mice subjected to pMCAO. **A**, Timeline of experimental procedure in adult (10-12 weeks) mice subjected to pMCAO. **B**, Representative Cresyl violet-stained coronal sections from vehicle- and adropin-treated mice at 21 days after pMCAO. **C**, Post-ischemic treatment with synthetic adropin^34-76^ peptide (900 nmol/kg; i.v.) at the onset of pMCAO significantly reduces brain infarct volume. Unpaired t-test, \*\*\**P*<0.001, n=9-10 per group. **D-L**, Neurobehavioral tests were performed on vehicle- and adropin-treated mice before (baseline) and on defined days (24h, 48h, 7d, 14d, and 21 d) after pMCAO as depicted in panel **A**. Sensorimotor function was evaluated by the adhesive removal test (**D, E**), cylinder test (**F, G**), open field test (**H-J**) and corner test (**K, L**). Recording data show that treatment with 900 nmol/kg of synthetic adropin peptide significantly reduces the time for mice to sense and remove the sticker on the affected paw (**D, E**). Also, mice treated with synthetic adropin show sustained recovery on impaired sensorimotor function, as shown by a dramatic increase in the percentage of first contact events with both forelimbs compared to the vehicle group (**G**). Graphical representation and representative tracking maps with corresponding heat maps of time spent per location in the open field chamber show that adropin-treated mice recovered better in total distance traveled at 48h after pMCAO than vehicle animals (**I, J**). In the corner test, a significant improvement on left turns (ipsilateral side) was observed at 48h post-stroke in mice treated with adropin compared to vehicle (**L**). **M**, There was no significant difference in body weight loss between adropin- and vehicle-treated mice over 21 days after pMCAO. Two-way ANOVA with Bonferroni post-tests, \**P*<0.05, \*\**P*<0.01, \*\*\**P*<0.001. Vehicle (n=10), Adropin (n=9).

### Transgenic overexpression of adropin reduces brain infarction while genetic adropin deficiency worsens ischemic brain injury in adult male and female mice

To further examine whether adropin signaling has neuroprotective effects in ischemic stroke, we compared the response of *Enho*^*-/-*^ mice (*18*) and AdrTg (*8*) to MCAO. We first characterized these two mouse strains to confirm adropin deficiency or overexpression compared to their respective littermate controls. Data clearly showed a single adropin band around 17 kDa in *Enho*^*+/+*^ mice but not in brain homogenates from the *Enho*^*-/-*^ mice (**Figures S3A and S3B**). Quantitative RT-PCR and western blot analysis confirmed that *Enho* gene expression and protein levels are significantly upregulated in the brain of AdrTg mice compared to the WT littermates (**Figure S3C through S3E**).

Homozygous deficiency of the *Enho* gene (*Enho*^*-/-*^) in male mice resulted in a significant increase in brain infarct volume compared to *Enho*^*+/+*^ controls, but there were no significant differences in infarct volume between *Enho*^*+/+*^ and *Enho*^*+/-*^ (heterozygous) mice after stroke (**Figure 3A and 3B**). Conversely, the global overexpression of adropin (AdrTg) in male mice resulted in a smaller brain infarction after 48h following pMCAO than the corresponding WT littermates (**Figure 3C and 3D**). This finding is in line with the pharmacological data using synthetic adropin peptide. Similarly, a larger infarct volume was observed in adult female *Enho*^*-/-*^ mice at 48h after pMCAO compared to *Enho*^*+/+*^ controls. In contrast, a smaller brain infarct size was observed in female AdrTg mice compared to their WT littermates (**Figure 3E through 3H**).

**Figure 3.**
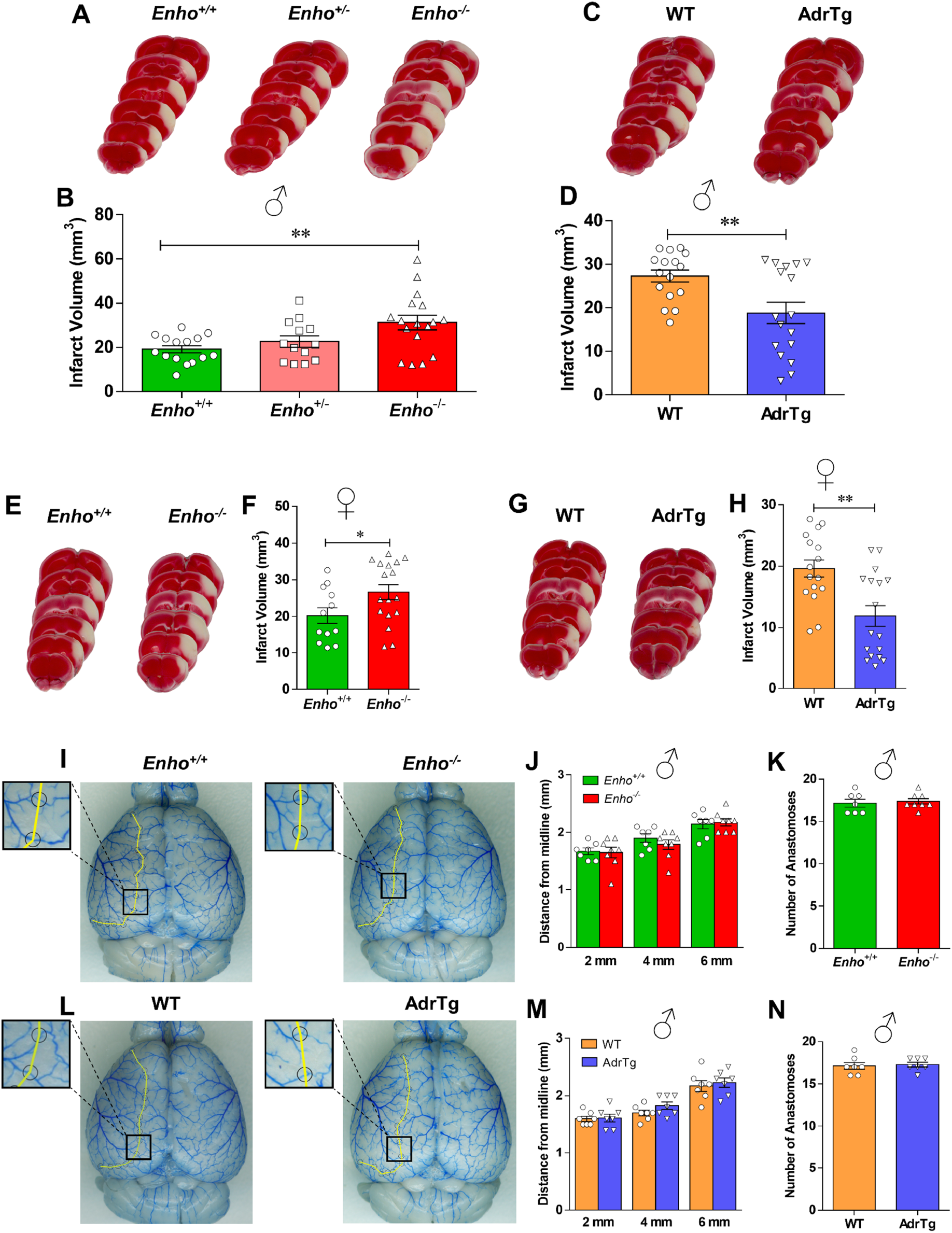
Levels of endogenous adropin determine the susceptibility to ischemic stroke injury in male and female mice with no significant differences in cerebrovascular anatomy among genotypes. **A-H**, Adult (10-12 weeks) male and female adropin overexpressing (AdrTg), heterozygous adropin-deficient (*Enho*^+/-^), homozygous adropin-deficient (*Enho*^-/-^) mice and their corresponding wild-type littermates were subjected to pMCAO and euthanized at 48h after cerebral ischemia to measure infarct volume by TTC staining. **A, B**, Representative images of TTC-stained brain sections and graphical data show that deficiency of *Enho* gene in male mice resulted in a significant increase in brain infarct volume compared to *Enho*^+/+^ controls. One-way ANOVA with Bonferroni post-tests, \*\**P*<0.01, n=13-17 per group. Conversely, transgenic overexpression of adropin (AdrTg) in male mice resulted in a smaller coronal brain infarction after 48h following pMCAO (**C**) and total infarct volume (**D**) compared to WT littermates. Unpaired t-test, \*\**P*<0.01, n=16-17 per group. Similarly, *Enho* deficiency in female mice also resulted in a significant increase in brain infarct volume compared to *Enho*^+/+^ controls (**E, F**). In contrast, a smaller brain infarct size was observed in female AdrTg mice at 48h after pMCAO compared to their WT littermates (**G, H**). Unpaired t-test, \**P*<0.05, \*\**P*<0.01, n=12-17 per group. **I-N**, Intravascular perfusion with latex blue shows permanent staining of small vessels on the dorsal surface of brains in adult (10-12 weeks) male *Enho*^-/-^, AdrTg and their corresponding wild-type littermates. A yellow line connected adjacent anastomosis points, and the distance from the midline to the line of anastomoses was measured at 2, 4, and 6 mm from the frontal pole. Quantification of the distance of the anastomotic points between the anterior cerebral artery (ACA) and the middle cerebral artery (MCA) from the midline showed no differences in the supplying territory of the MCA between *Enho*^-/-^ and *Enho*^+/+^ mice (**I-K**), as well as the AdrTg and the corresponding WT mice (**L-N**). Unpaired t-test. *Enho*^+/+^ (n=7), *Enho*^-/-^ (n=8), WT (n=7), AdrTg (n=7).

To rule out the possibility that changes in major physiological parameters could explain the differential vulnerability to ischemic stroke damage among the different mouse strains, arterial blood pH, blood oxygen (PO_2_), carbon dioxide saturation (PCO_2_), ion concentrations, blood glucose, hematocrit, hemoglobin, blood pressure, and heart rate were measured in adult male *Enho*^-/-^, AdrTg and their corresponding wild-type littermates. As shown in **Table S1**, no significant differences in these physiological parameters were observed between *Enho*^*-/-*^ and *Enho*^*+/+*^ mice or between AdrTg and the corresponding WT control mice.

Strain-dependent variability in cerebral vascular anatomy might be associated with different susceptibility to cerebral ischemia (*19*). We analyzed the vasculature on the dorsal surface of the brain and large vessels on the ventral surface of the brain in adult male *Enho*^-/-^, AdrTg and their corresponding wild-type littermates. Morphological analysis of cerebral vasculature and anastomoses using latex blue vessel casting indicated no significant effects of genotype on the distance of the anastomotic points between the anterior cerebral artery (ACA) and the middle cerebral artery (MCA) from the midline in the supplying territory of the MCA in *Enho*^*-/-*^ mice (**Figure 3I** through **3K**) or AdrTg mice (**Figure 3L** through **3N**). No significant differences were observed in the diameters of the ACA, MCA, internal carotid artery (ICA), posterior communicating artery (PCA), and basilar artery (BA) in the different adropin mouse strains (**Figure S4A** through **S4D**). Changes in either physiological variables or cerebral vascular anatomy are unlikely to contribute to the different stroke outcomes among the mouse strains utilized in this study.

### Treatment with synthetic adropin peptide^34-76^ prevents stroke-induced loss of endogenous adropin, reduces neurovascular injury, MMP-9 activity, oxidative damage, and neutrophil infiltration

Adropin has been implicated in preserving endothelial barrier integrity *in vitro* and *in vivo* (*5-7*). We, therefore, quantified the extravasation of plasma proteins into the ischemic brain as a measure of blood-brain barrier (BBB) injury in vehicle- and adropin-treated mice subjected to pMCAO. At 24h after stroke, we found a dramatic increase in immunoglobulin G (IgG), albumin, and hemoglobin (Hb) in the ipsilateral cortex compared to the contralateral side of ischemic animals and sham controls; post-ischemic treatment with synthetic adropin peptide significantly reduced the observed increased levels of these markers (**Figure 4A** through **4C**). In agreement with these findings, adropin-deficiency significantly increased the extravasation of IgG and albumin into the ipsilateral cortex compared to *Enho*^*+/+*^ mice. In contrast, AdrTg exhibited significantly reduced leakage of IgG and albumin into the ischemic brain (**Figure S5A** through **S5D**). Additional findings revealed that adropin treatment prevented stroke-induced degradation of tight junction proteins (TJPs), as seen by preservation of zona occludens (ZO)-1 and occludin levels (**Figure 4D** through **4F**).

**Figure 4.**
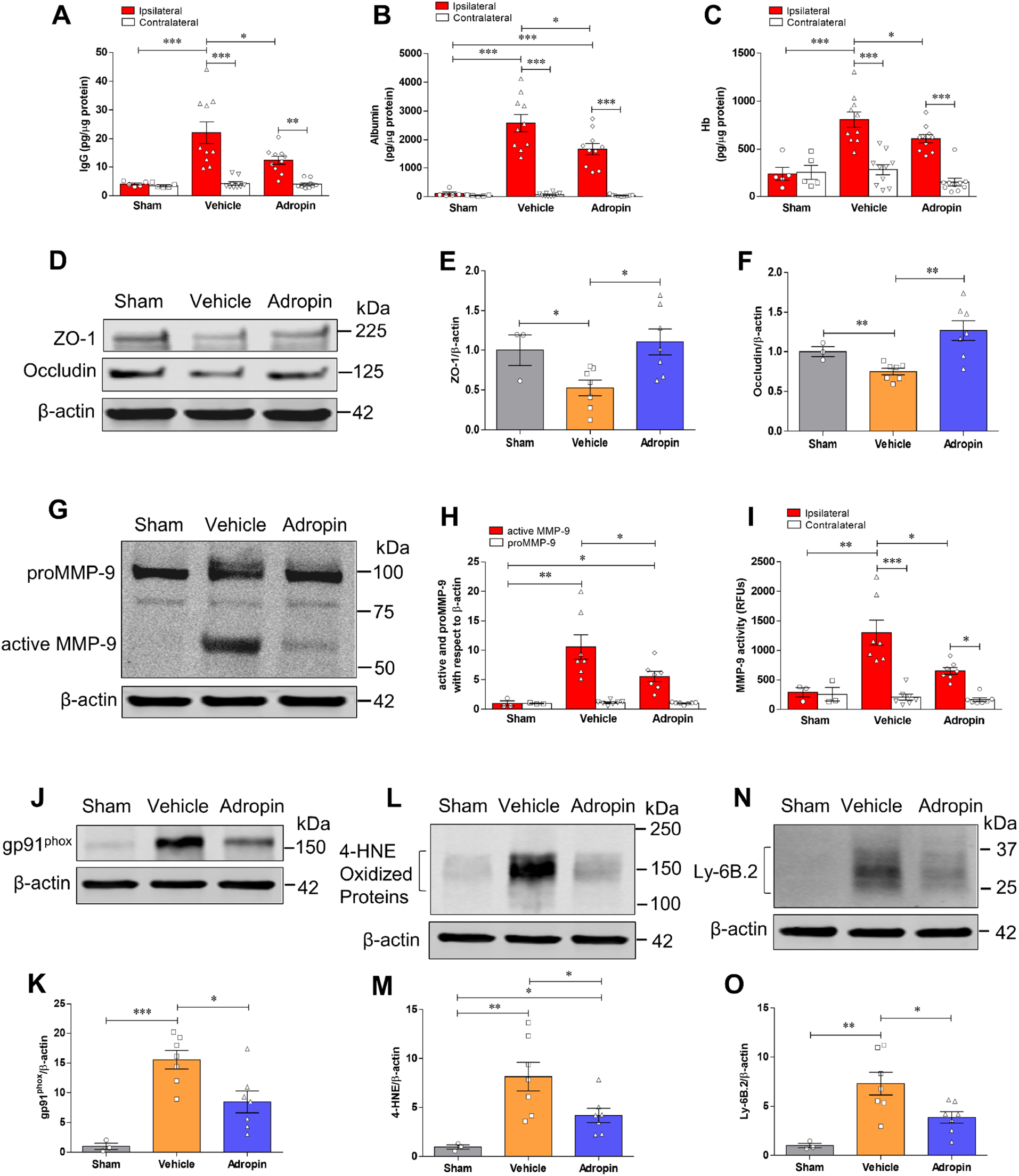
Post-ischemic treatment with synthetic adropin peptide reduces neurovascular injury, MMP-9 activity, oxidative damage, and neutrophil infiltration in the ischemic brain. Adult (10-12 weeks) male mice received one bolus injection of vehicle or synthetic adropin^34-76^ peptide (900 nmol/kg; i.v.) at the onset of ischemia and were euthanized 24h after pMCAO and cerebral cortex from ipsilateral and contralateral sides were collected for molecular analysis. Sham-operated mice received the same surgical procedures except for the MCA occlusion. **A-C**, Stroke induced a dramatic BBB damage, which was assessed by increased levels of IgG and albumin as well as hemorrhagic transformation quantified by increased hemoglobin level in the ipsilateral cortex, and post-ischemic treatment with adropin significantly reduced the increased levels of these markers. Two-way ANOVA with Bonferroni post-tests, \**P*<0.05, \*\**P*<0.01, \*\*\**P*<0.001. Sham (n=5), Vehicle (n=10), Adropin (n=10). **D**, Representative immunoblots for tight junction proteins ZO-1 and occludin in homogenates from the ischemic cortex. β-actin was used as a loading control. **E, F**, Densitometric analysis shows that stroke resulted in significant degradation of ZO-1 and occludin in the ischemic cortex. The degradation of these two tight junction proteins was dramatically attenuated in animals receiving adropin treatment. One-way ANOVA with Bonferroni post-tests, \**P*<0.05, \*\**P*<0.01. Sham (n=3), Vehicle (n=7), Adropin (n=7). **G-I**, Post-ischemic treatment with adropin significantly reduced MMP-9 levels measured by Western blot (**G, H**) and enzymatic activity by immunocapture assay (**I**) in the ipsilateral cerebral cortex compared to the vehicle group. RFUs: relative fluorescence units. One-way ANOVA with Bonferroni post-tests, \**P*<0.05, \*\**P*<0.01, \*\*\**P*<0.001. Sham (n=3), Vehicle (n=7), Adropin (n=7). **J-O**, Representative Western blots, and densitometric analysis showed that stroke induced a dramatic increase in oxidative stress markers, NADPH oxidase isoform NOX2 (gp91^phox^) and 4-hydroxy-2-nonenal (4-HNE)-modified proteins, as well as infiltration of neutrophils (Ly-6B.2 as a specific neutrophil marker) into the ischemic cerebral cortex. Post-ischemic treatment with adropin significantly reduced the increased levels of these oxidative stress and inflammatory markers. One-way ANOVA with Bonferroni post-tests, \**P*<0.05, \*\**P*<0.01, \*\*\**P*<0.001. Sham (n=3), Vehicle (n=7), Adropin (n=7).

In an initial effort to identify possible mechanisms underlying adropin’s neurovascular protection, we quantified MMP-9, gp91^phox^-containing NADPH oxidase (NOX2), as well as levels of 4-hydroxy-2-nonenal (4-HNE)-modified proteins in the ischemic brain of mice given vehicle or adropin (900 nmol/kg; i.v.) at the start of pMCAO. Increased MMP-9 and oxidative stress are critical mediators of BBB breakdown in ischemic stroke (*20-22*). Adropin treatment significantly reduced both active MMP-9 protein levels (**Figure 4G and 4H**) and MMP-9 enzymatic activity (**Figure 4I**). There was a marked increase in gp91^phox^ levels and 4-HNE-modified proteins in the ischemic cerebral cortex at 24h after pMCAO, and adropin treatment significantly reduced gp91^phox^ and protein oxidation (**Figure 4J** through **4M**). As infiltrating neutrophils are the primary source of active MMP-9 in the brain following stroke (*23*), we measured the levels of Ly-6B.2, a neutrophil marker, in the ischemic cerebral cortex at 24h after pMCAO. We found a dramatic infiltration of neutrophils into the ischemic cortex of vehicle-treated mice, and this increase was significantly attenuated in the adropin-treated group (**Figure 4N and 4O**). Adropin thus plays an essential role in the preservation of BBB integrity in ischemic stroke.

As anticipated, pMCAO suppressed endogenous adropin levels in the ischemic cerebral cortex at 24h, and this loss can be significantly attenuated by the treatment with synthetic adropin^34-76^ (**Figure S6A and S6B**). As shown in **Figure S6C**, the immunoblotting image clearly indicates that the molecular weight of the endogenous adropin in the mouse brain is around 17 kDa. In comparison, the synthetic adropin^34-76^ peptide is about 14 kDa, suggesting that endogenous adropin is significantly reduced after ischemic stroke. We found that endogenous brain adropin is likely a membrane-bound protein, since extraction of proteins without a detergent dramatically reduced the adropin signal detected by immunoblotting (**Figure S6D**), confirming previous data that adropin is a membrane-bound protein highly expressed in the brain (*9*).

### Neuroprotective effects of adropin in ischemic stroke are mediated by an eNOS-dependent mechanism

Lovren *et al*. demonstrated that adropin reduced endothelial permeability by increasing eNOS phosphorylation at serine 1177 (Ser^1177^) (*5*). To determine the role of eNOS in adropin-mediated neuroprotection in ischemic stroke, we examined the effects of adropin treatment on eNOS phosphorylation at Ser^1176^ following pMCAO (mouse phosphorylation site corresponds to Ser^1177^ in human eNOS). We found an increase in eNOS phosphorylation in the ischemic cortex at 24h after pMCAO compared with sham, and stroke mice treated with adropin had a dramatic increase in eNOS phosphorylation level, which were significantly higher than vehicle and sham controls (**Figure 5A and 5B**). The findings that adropin can increase eNOS phosphorylation at Ser^1176^, a critical post-translational modification indispensable for its enzymatic activity (*24*), prompted us to investigate whether adropin treatment affects infarct size 48h after pMCAO in adult male eNOS knockout (*eNOS*^*-/-*^) mice. While adropin (900 nmol/kg; i.v. at the onset of pMCAO) was remarkably protective in wild-type (*eNOS*^*+/+*^) control mice, it failed to confer protection in *eNOS*^*-/-*^ mice (**Figure 5C and 5D**). This result strongly suggests that signaling through eNOS is required for the protection by adropin against ischemic stroke injury.

**Figure 5.**
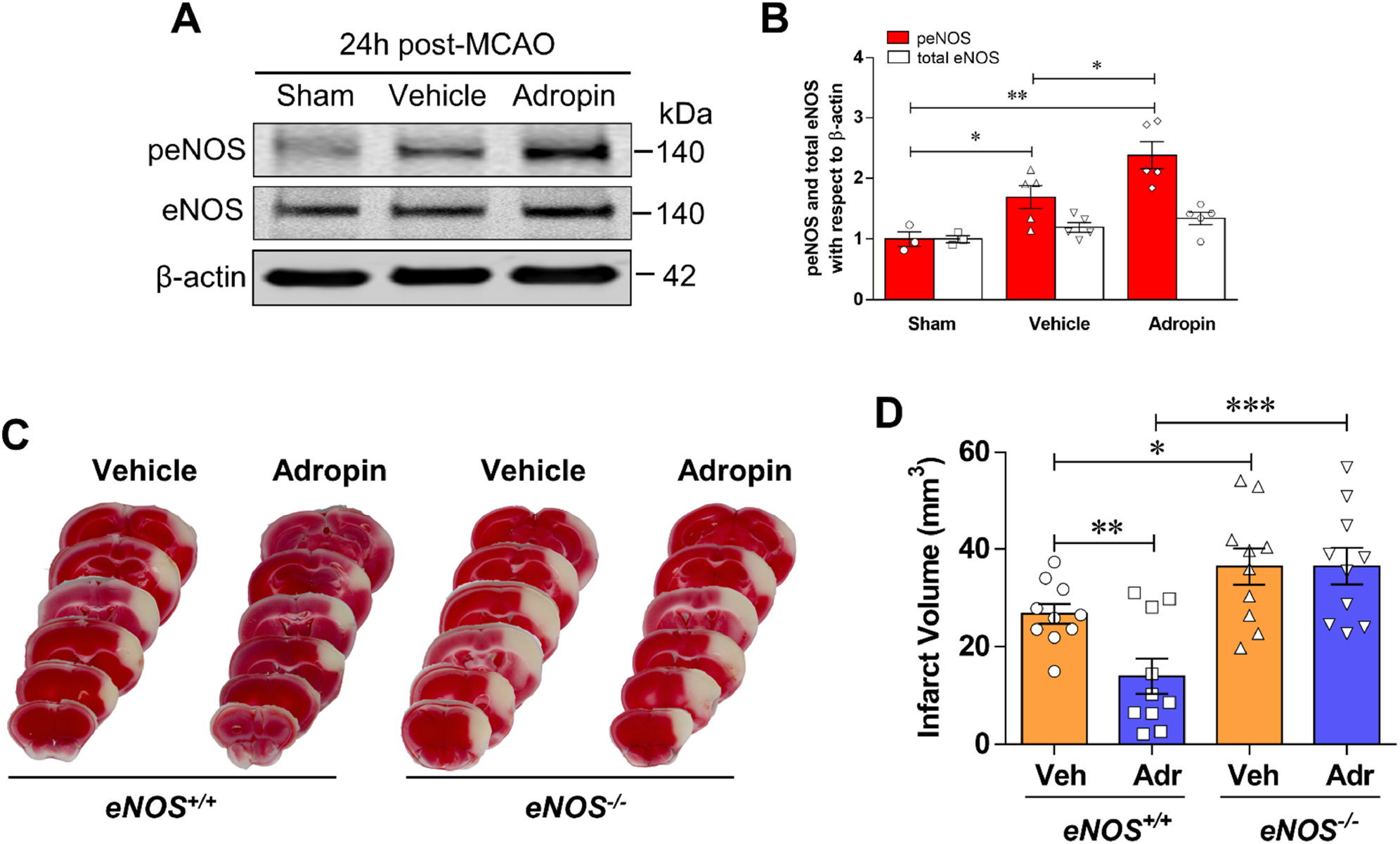
Neuroprotective effects of adropin are dependent on eNOS. **A, B**, Adult (10-12 weeks) male mice were given one dose of either vehicle or synthetic adropin^34-76^ peptide (900 nmol/kg; i.v.) at the onset of cerebral ischemia and euthanized at 24h post-stroke. Phospho- and total-eNOS levels in cortical homogenates were quantified by immunoblotting. Representative Western blots and densitometric analysis showed that adropin treatment significantly increased eNOS phosphorylation at Ser^1176^ following pMCAO. One-way ANOVA with Bonferroni post-tests, \**P*<0.05, \*\**P*<0.01. Sham (n=3), Vehicle (n=5), Adropin (n=5). **C, D**, Mice received vehicle or synthetic adropin^34-76^ peptide (900 nmol/kg; i.v.) at the onset of pMCAO and euthanized at 48h. Representative photographs of coronal brain sections stained with TTC in *eNOS*^+/+^ and *eNOS*^-/-^ mice with and without adropin treatment (**C**). Adropin treatment resulted in a significant decrease in brain infarct volume compared with the vehicle group in *eNOS*^+/+^ mice. These protective effects of adropin on infarct volume were completely abolished in *eNOS*^-/-^ mice (**D**). One-way ANOVA with Bonferroni post-tests, \**P*<0.05, \*\**P*<0.01, \*\*\**P*<0.001, n=10 per group.

As activation of eNOS affects blood flow, we next examined whether adropin treatment changes cerebral blood flow (CBF) in mice. Compared to the vehicle group, mice that received a bolus injection of adropin^34-76^ (900 nmol/kg) or acetylcholine (ACh, 1.0 mM; used as a comparison) via the tail vein had a significant increase in cortical CBF up to 30 min following the intravenous injection (**Figure 6A through 6C**). Adropin peptide shows more potent effects on CBF than ACh treatment. These data are the first to demonstrate that adropin increases CBF, which further supports the important role of this peptide in the regulation of endothelial function (*5, 6, 10, 13*). To further explore the underlying mechanisms of eNOS activation by adropin, we measured phosphorylation levels of Akt and ERK1/2, two upstream effectors of eNOS phosphorylation, as well as quantified levels of eNOS phosphorylation in the brain of mice treated with synthetic adropin peptide or in the AdrTg mice. There is increasing evidence showing that PI3K/Akt and ERK1/2 signaling mechanisms mediate the effects of adropin *in vitro* and *in vivo* (*5, 25-27*). To investigate whether adropin increased the phosphorylated protein levels of eNOS, Akt, and ERK1/2 in brain endothelial vessels, we extracted brain microvessels from adult male mice at 30 min after intravenous injection of vehicle or 900 nmol/kg synthetic adropin peptide. Quality and purity of brain microvessels were assessed by microscopy and western blots using endothelial cell and tight junction protein markers, indicating that the brain microvessel fraction was enriched compared to total homogenates (**Figure 6D**). As shown in **Figure 6E through 6H**, adropin treatment significantly increased the phosphorylated protein levels of eNOS, Akt, and ERK1/2. Notably, plasma levels of nitrite/nitrate (NO metabolites) were significantly increased at 30 min following synthetic adropin peptide injection (**Figure 6I**), likely suggesting increased eNOS activity. In line with these findings, transgenic overexpression of adropin also significantly upregulated the phosphorylated protein levels of eNOS, Akt, and ERK1/2 in the cerebral cortex (**Figure 6J** through **6N**). Collectively, these data demonstrate that treatment with exogenous adropin peptide or overexpression of endogenous adropin induces eNOS phosphorylation associated with increased phosphorylated protein levels of Akt and ERK1/2, which likely contributes to the neurovascular protection by adropin in ischemic stroke.

**Figure 6.**
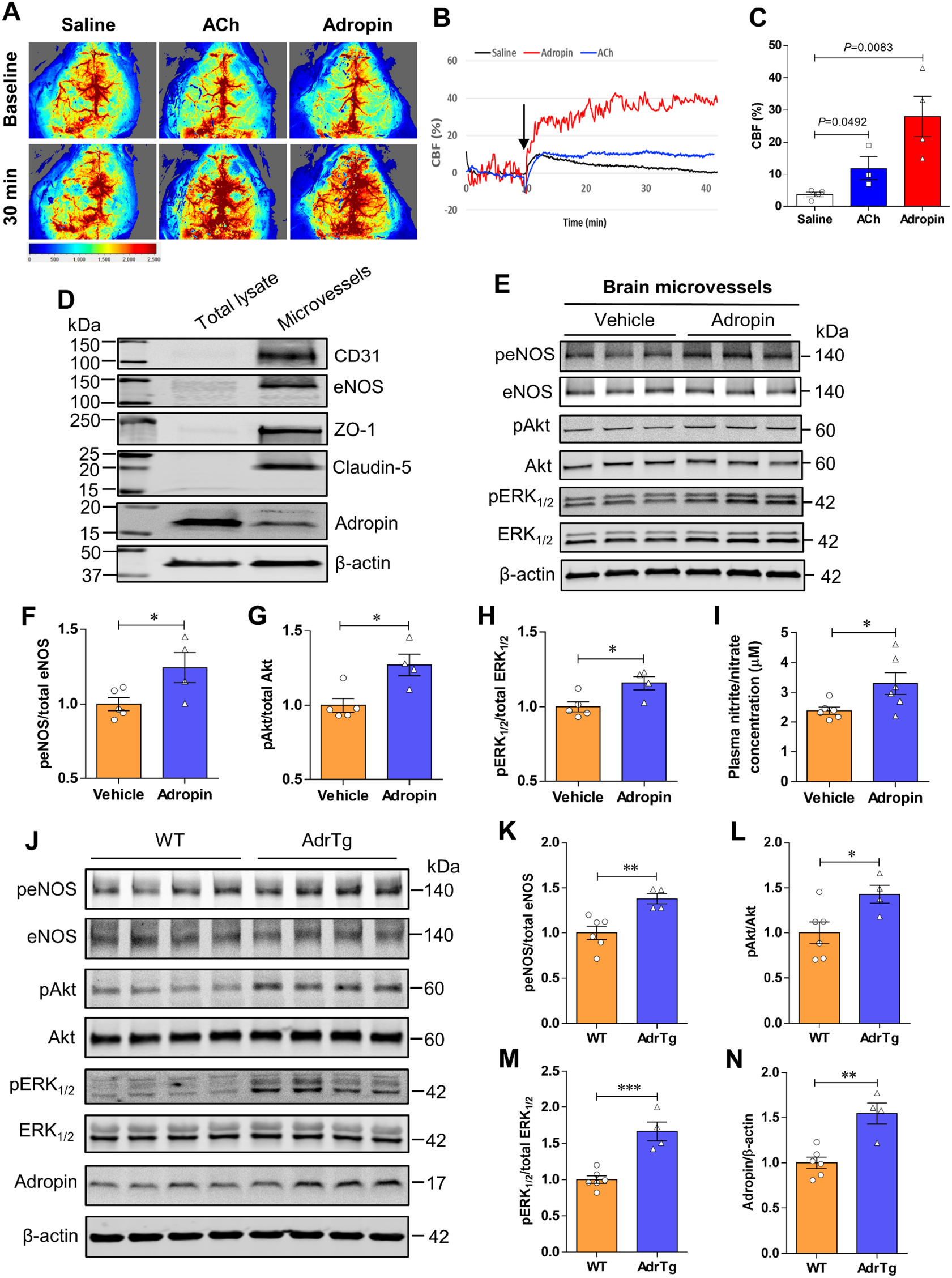
Adropin increases phosphorylated protein levels of eNOS, Akt, and ERK1/2 in the mouse brain. **A-C**, Adult (10-12 weeks) male mice received one bolus injection of vehicle, synthetic adropin^34-76^ peptide (900 nmol/kg), or acetylcholine (ACh, 1.0 mM) via the tail vein, and cerebral blood flow (CBF) was recorded with the MoorFLPI software. The average CBF change (CBF %) within a total recording time of 30 min was normalized to the baseline CBF (first 10 min). Representative image and recording of CBF measured by laser speckle flowmetry at different time points (**A, B**). Statistical analysis of CBF shows that both adropin and ACh (used for comparison) significantly increased cortical CBF compared to the vehicle group (**C**). Unpaired t-test, n=4 per group. **D-I**, Adult male mice received one bolus injection of vehicle or synthetic adropin^34-76^ peptide (900 nmol/kg; i.v.) and were euthanized at 30 min for brain microvessel isolation as described in detail in the Methods section. **D**, Twenty micrograms of protein from brain total lysates or microvessels were loaded to a 4-20% SDS-polyacrylamide gel, and several targeted proteins were detected by immunoblotting. Representative western blots show that endothelial cell markers such as CD31, eNOS, ZO-1, and Claudin-5 and cytoskeletal protein β-actin are enriched in the brain microvessel fraction. Adropin shows a higher level in the brain lysate fraction. Data are representative of three independent experiments. **E**, Representative western blots for phosphorylated and total eNOS, Akt, and ERK1/2 in homogenates from brain microvessels with β-actin as a loading control. **F-H**, Densitometric analysis shows that treatment with synthetic adropin significantly increased the phosphorylated protein levels of eNOS, Akt, and ERK1/2. Unpaired t-test, \**P*<0.05. Vehicle (n=5), Adropin (n=4). **I**, Plasma levels of nitrite/nitrate (NO metabolites) in vehicle- and adropin-treated mice at 30 min after administration were measured with a nitrite/nitrate colorimetric assay. Graphical data show that adropin treatment significantly increased NO metabolite levels in plasma compared to the vehicle group. Unpaired t-test, \**P*<0.05. Vehicle (n=6), Adropin (n=6). **J**, Representative western blots for phosphorylated and total eNOS, Akt and ERK1/2, and adropin in homogenates from brain cortex in naïve adult male WT and adropin transgenic (AdrTg) mice with β-actin as a loading control. **K-N**, Densitometric analysis shows that protein levels of phospho-eNOS, -Akt, and -ERK1/2 and adropin were significantly higher in the AdrTg mice than WT controls. Unpaired t-test, \**P*<0.05, \*\**P*<0.01, \*\*\**P*<0.001. WT (n=6), AdrTg (n=4).

### Adropin reduces infarct volume and improves long-term neurobehavioral outcomes in aged male mice following stroke

Advanced age is the most critical risk factor for stroke (*28*). Limiting preclinical studies to young, healthy mice is an important reason for failing to translate basic studies to the clinic (*29*). We, therefore, investigated whether adropin confers long-lasting neuroprotection in aged mice subjected to ischemic stroke. As shown in **Figure 7A**, a battery of neurobehavioral tests assessing sensorimotor and recognition memory functions were performed in aged (18-24 months) male mice with genetic overexpression or deficiency of adropin subjected to pMCAO. MRI-based infarct size analysis was performed at 14d following stroke. *Enho*^*-/-*^ mice displayed a significant increase in brain infarct volume compared to *Enho*^*+/+*^ controls (**Figure 7B and 7C**), while AdrTg mice had a smaller brain infarction than WT littermates (**Figure 7D and 7E**). Sensorimotor function assessed by the adhesive removal test was significantly impaired up to 14 days after pMCAO in *Enho*^*-/-*^ mice compared with *Enho*^*+/+*^ mice (**Figure 7F and 7G**). There was no significant difference in body weight loss between *Enho*^*+/+*^ and *Enho*^*-/-*^ mice (**Figure 7J**). However, the sensorimotor deficits were significantly alleviated at 7 and 14d after pMCAO in AdrTg mice compared with WT mice (**Figure 7H and 7I**). AdrTg mice experienced less body weight loss (**Figure 7K**). Unexpectedly, there was no significant difference observed in nesting behavior between *Enho*^*+/+*^ and *Enho*^*-/-*^ and WT and AdrTg mice. However, the nest-building ability was impaired in these four strains at 48h after pMCAO (**Figure S7A through S7F**). Further, AdrTg mice had much better recovery in NOR at 14d after pMCAO than the WT mice. *Enho*^*-/-*^ mice displayed worse recognition memory deficits when compared to the *Enho*^*+/+*^ controls after stroke (**Figure 7L through 7P**). These data suggest that endogenous adropin exerts beneficial effects on reducing infarct volume and improving long-term neurological deficits in aged mice subjected to ischemic stroke injury.

**Figure 7.**
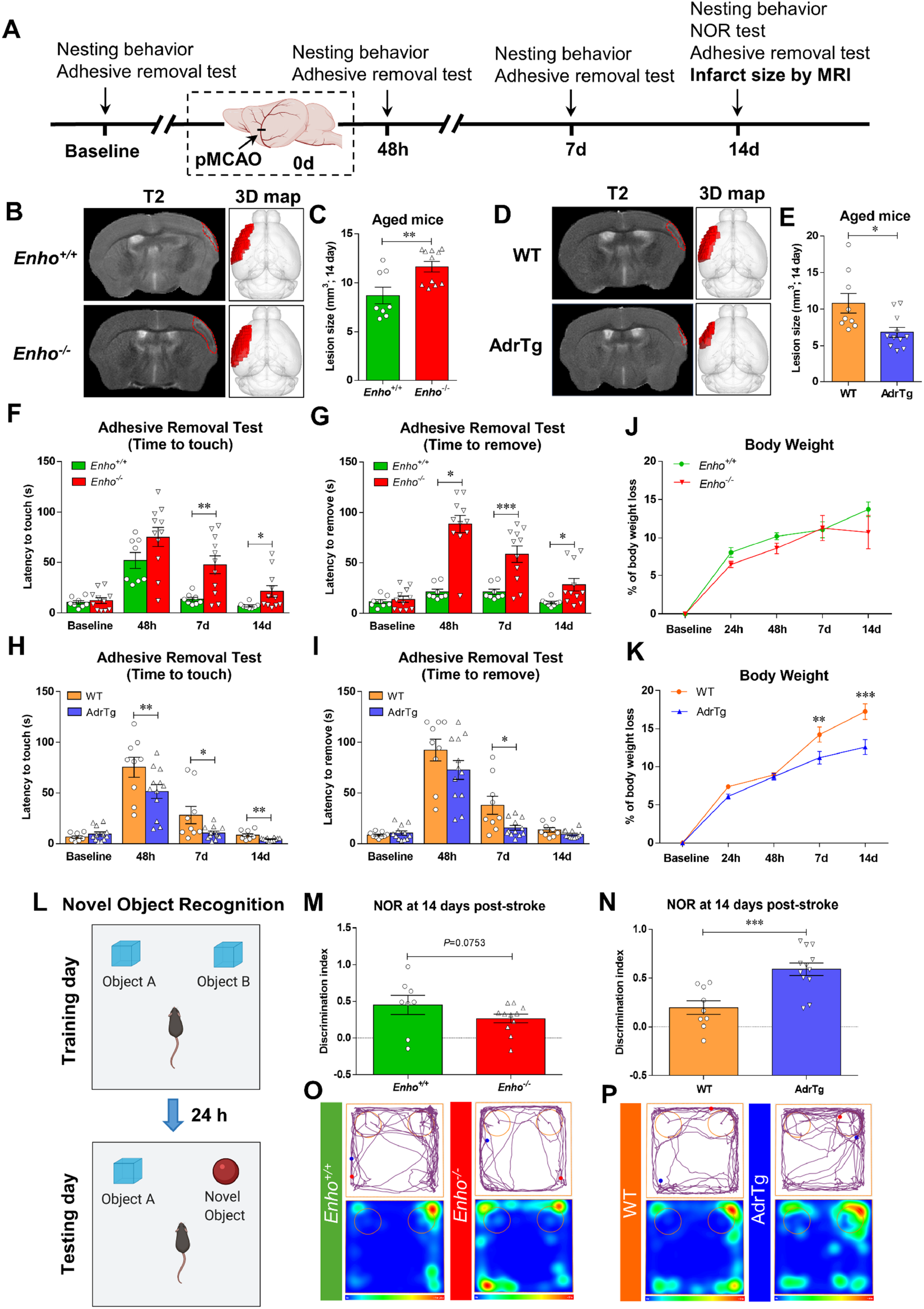
Effects of endogenous adropin on infarct volume and long-term neurobehavioral outcomes in aged mice subjected to pMCAO. **A**, Timeline of behavioral testing and magnetic resonance imaging (MRI) scanning in aged (18-24 months) male mice subjected to pMCAO. **B-E**, Representative T2-weighted MRI scans and 3-dimensional (3D) reconstruction images of the ischemic lesion in adropin knockouts (*Enho*^-/-^), adropin overexpressing mice (AdrTg) and their corresponding wild-type littermates at 14 days after pMCAO (**B, D, *Left:*** T2-MRI for *Enho*^+/+^ vs. *Enho*^-/-^, WT vs. AdrTg; ***Right*:** 3D-image for *Enho*^+/+^ vs. *Enho*^-/-^, WT vs. AdrTg). The ischemic area was marked with a discontinuous red line in T2 maps and depicted as a dark red area in the left hemisphere in 3D images. Graphical data show that deficiency of the *Enho* gene resulted in a significant increase in brain infarct volume compared to *Enho*^+/+^ controls (**C**). In contrast, transgenic overexpression of adropin resulted in a smaller brain infarction than WT littermates (**E**). Unpaired t-test, \**P*<0.05, \*\**P*<0.01. *Enho*^+/+^ (n=8), *Enho*^-/-^ (n=11), WT (n=9), AdrTg (n=11). **F-P**, Neurobehavioral tests were performed before (baseline) and defined days (48h, 7d, and 14d) after pMCAO as depicted in panel **A**. Sensorimotor and recognition memory functions were evaluated by adhesive removal test (**F, G, H, I**) and novel object recognition (**L-P**). Deficiency of the *Enho* gene significantly increased the time mice took to sense and remove the sticker on the affected paw at days 7 and 14 after stroke compared to *Enho*^+/+^ mice (**F, G**). In contrast, AdrTg mice took significantly less time to sense and remove the sticker on the affected paw at day 7 than the corresponding WT littermates (**H, I**). Graphical representation, tracking maps, and corresponding heat maps of time spent per location in the open field chamber show that *Enho*^-/-^ mice have a worse recovery in long-term recognition memory performance at 14d after pMCAO compared to the *Enho*^+/+^ mice (**M, O**). Still, AdrTg mice show much better cognitive function recovery than the WT littermates (**N, P**). There is no significant difference in body weight loss between *Enho*^+/+^ and *Enho*^-/-^ mice over 14 days after stroke (**J**). Bodyweight loss was significantly attenuated after 48 hours following pMCAO in the AdrTg mice compared to the WT littermates (**K**). Two-way ANOVA with Bonferroni post-tests, \**P*<0.05, \*\**P*<0.01, \*\*\**P*<0.001. *Enho*^+/+^ (n=8), *Enho*^-/-^ (n=11), WT (n=9), AdrTg (n=12).

## Discussion

Ischemic stroke remains a devastating neurological disorder with minimal treatment options. Reperfusion therapies utilizing mechanical thrombectomy or tPA are the only clinical options for a limited group of stroke patients who qualify to receive them. Ischemic brain injury progresses over time, even in patients that have been successfully recanalized. Thus, there is an urgent need to identify effective neuroprotective agents to minimize cell death and improve functional outcomes after stroke. The present study clearly demonstrates that post-ischemic treatment with synthetic adropin peptide or transgenic overexpression of endogenous adropin exerts beneficial effects in a preclinical model of ischemic stroke. Notably, adropin deficiency worsens stroke outcomes, suggesting that endogenous adropin plays a critical role in modulating the brain’s susceptibility to ischemia. From a translational point of view, findings that treatment with synthetic adropin reduces infarct volume and improves long-term neurological recovery in both young and aged mice are of great significance. The adropin peptide sequence could provide the foundation for developing a novel biologic therapy administered after injury to confer neuroprotection to the ischemic brain.

Mechanistically, we found that adropin treatment reduces stroke-induced BBB damage, MMP-9 activity, oxidative damage, and infiltration of neutrophils into the ischemic brain. Adropin’s neuroprotective effects are dependent on eNOS. Additionally, adropin^34-76^ treatment significantly increases CBF, an effect associated with increased levels of phosphorylated eNOS, Akt, and ERK1/2 in brain microvessels and elevated nitrite/nitrate levels in plasma. Consistently, transgenic overexpression of endogenous adropin also significantly increases phosphorylated eNOS, Akt, and ERK1/2 in the cerebral cortex in the naïve mouse brain. Our current findings are the first to demonstrate that adropin is neuroprotective in stroke, likely by reducing BBB damage and improving cerebral perfusion through eNOS-dependent signaling mechanisms.

Since knowledge of the cellular distribution of adropin in the brain is limited, we determined the cell types expressing this peptide in the naïve mouse brain. We used double immunostaining with a highly specific adropin monoclonal antibody or an adropin-internal ribosomal entry sequence (IRES)-Cre-TdTomato reporter mouse (*14*). We consistently found that adropin is highly expressed in brain endothelial cells and neurons, with minimal expression in astrocytes or microglial cells. These findings partly differ from a previous report using an adropin polyclonal antibody showing that adropin is expressed in several cell types in the rat brain, including endothelial cells, pericytes, and astrocytes (*7*). Both studies consistently indicate that adropin is expressed in the brain vasculature. Here, we show that adropin is abundantly expressed in highly-vascularized organs, including brain, liver, kidney, and lung, moderately expressed in spleen and intestines, and poorly expressed in heart and skeletal muscle, which is in agreement with the findings that adropin immunoreactivity is primarily observed in the vascular or peritubular area of brain, cerebellum, and kidney (*30*). Taken together, these data suggest that adropin plays an essential role in regulating endothelial function in the brain and other organs.

There is increasing clinical evidence that low serum adropin levels correlate with endothelial dysfunction in diabetes and coronary artery disease patients (*11, 31, 32*). Our findings are the first to reveal that adropin levels are significantly reduced in the ischemic brain from 4 to 48 hours, which is associated with a decrease in plasma adropin at 48h in mice following ischemic stroke. These results align with findings from a recent report showing lower serum adropin in patients with acute ischemic stroke (*17*). However, this differs from another study where transient MCAO does not change plasma adropin levels in streptozotocin-induced diabetic rats. In contrast, plasma adropin is significantly increased in non-diabetic rats following a stroke compared to sham-operated control animals (*33*). The apparent discrepancies in stroke-induced changes in circulating adropin between these studies could be due to differences in stroke models, injury severity, species, and the use of different immunoassays to quantify adropin in plasma. More studies are needed to investigate the impact of stroke on circulating adropin levels and how this correlates with stroke outcomes.

Loss of endogenous adropin in the ischemic brain was associated with stroke injury, and adropin peptide treatment can reduce stroke damage and restore brain adropin level, suggesting that adropin plays an important role in ischemic stroke pathology. We found that adropin treatment provides robust improvement of sensorimotor function in stroke mice. Importantly, we found that the protective effects of adropin are also observed in aged mice subjected to ischemic stroke and that adropin overexpressing mice show better recognition memory performance, which agrees with our recent observation that adropin treatment or adropin transgenic overexpression can significantly improve cognitive function in aged mice (*14*). Taken together, our data suggest that maintenance of brain adropin level could be pivotal for mitigating ischemic brain damage and improve long-term functional outcomes.

Focal cerebral ischemia is caused by a sudden interruption of cerebral blood flow resulting in mitochondrial dysfunction, energy failure, loss of ionic homeostasis, which subsequently leads to several injury cascades, including endothelial dysfunction and oxidative stress, where free radicals such as nitric oxide (NO) and superoxide (O_2_^-.^) are thought to be critical mediators (*34*). It is well-documented that endothelial nitric oxide synthase (eNOS)-derived NO exerts multiple pleiotropic functions, including modulation of blood flow, angiogenesis, and neurogenesis in ischemic stroke (*35*). Other studies have shown that eNOS is protective in ischemic stroke (*36-39*), and eNOS phosphorylation at Ser^1176^ (based on mouse eNOS sequence and equivalent to human eNOS at Ser^1177^ or bovine eNOS at Ser^1179^) is a key post-translational modification indispensable for its enzymatic activity (*24*). Importantly, deficient phosphorylation of eNOS at Ser^1176^, a hallmark of endothelial dysfunction, is associated with worse outcomes following stroke (*37, 38*). Increased eNOS activity is protective in the context of brain ischemia (*37, 39*), and eNOS-deficient mice have a worse stroke outcome (*36*). The bioavailability of eNOS-derived NO can be dramatically reduced due to increased O_2_^-.^ generation from NADPH oxidase (NOX), mainly the NOX2 isoform (also known as gp91^phox^). The reaction between NO and O_2_^-.^ forms peroxynitrite (ONOO^-^) (*40*), which is a highly reactive species known to activate matrix metalloproteinase (MMP)-9 and exacerbate BBB disruption and neuronal death after stroke (*41*).

Lovren *et al*. were the first to report that adropin regulates the endothelial function and promotes cell proliferation and angiogenesis (*5*). This effect is possibly mediated by upregulating VEGFR2 and its downstream effectors PI3K/Akt and ERK1/2 pathways, and thus activating eNOS to increase NO production in human endothelial cells *in vitro* (*5*). We previously showed that loss of adropin in aged rat brains correlates with reduced levels of eNOS and increased oxidative stress (*13*). A very recent study demonstrated that aerobic exercise training restores the reduced levels of adropin in aortic tissue and plasma in aged mice, which are associated with increased phosphorylation of Akt and eNOS (*10*). In line with these findings, the present study shows that post-ischemic treatment with adropin peptide dramatically increases eNOS phosphorylation at the Ser^1176^ site in the ischemic brain compared to sham or vehicle-treated mice. Interestingly, the protective effects of adropin on infarct volume are completely abolished in eNOS-deficient mice subjected to ischemic stroke. As expected, a significant increase in CBF was also observed in adropin-treated mice, which was associated with increased levels of eNOS phosphorylation in brain microvessels and nitrite/nitrate in plasma. Also, eNOS phosphorylation is increased in the brain of adropin transgenic mice. Collectively, these data strongly suggest that eNOS is involved in the adropin-mediated neuroprotection in ischemic stroke.

Emerging evidence from *in vitro* studies shows that adropin can increase the phosphorylation of Akt at Ser^473^ site (*5, 25*) and ERK1/2 at Thr^202^/Tyr^204^ sites (*5, 25, 42*), whereas pharmacological inhibition of PI3K/Akt with LY294002 and ERK1/2 with PD98059 abolished the protective effects of adropin in H9c2 cardiomyoblast cells (*25, 42*). Also, mice receiving adropin plasmid exhibited improved limb perfusion following hindlimb ischemia which was associated with higher eNOS, Akt, and ERK1/2 phosphorylation in gastrocnemius muscle relative to null plasmid-treated animals (*5*). As the eNOS upstream effectors, Akt or ERK1/2 can phosphorylate eNOS at Ser^1176,^ thus resulting in NO release (*43*). Lovren *et al*. demonstrated that either inhibition of PI3K/Akt with LY294002 or ERK1/2 with PD98059 could altogether abolish adropin-induced eNOS phosphorylation in human umbilical vein endothelial cells (HUVECs) (*5*). Genetic deletion of adropin decreased both Akt and eNOS phosphorylation levels in the lungs of *Enho*^*-/-*^ mice (*26*). Here, we found for the first time that adropin treatment induces eNOS phosphorylation associated with increased levels of phosphorylated Akt and ERK1/2 in brain microvessels. Moreover, we show that genetic overexpression of adropin also consistently upregulates eNOS, Akt, and ERK1/2 phosphorylation in the mouse brain. Taken together, this evidence suggests that Akt- and ERK1/2-dependent eNOS signaling pathways are likely involved in the adropin-mediated neuroprotection in ischemic stroke.

It is well documented that stroke-induced oxidative stress could impair eNOS-derived NO bioavailability, thus resulting in endothelial dysfunction and worse stroke outcomes (*44*). Adropin prevents nonalcoholic fatty liver disease through regulating oxidative stress mediators in the liver (*45, 46*). In a palmitate-induced liver fibrosis model, a gradient decrease in *Enho* mRNA expression was observed in primary murine hepatocytes treated with palmitate in a dose-dependent manner, which was associated with increased oxidative stress levels, including intracellular reactive oxygen species (ROS), hydrogen peroxide (H_2_O_2_), superoxide (O_2_^-.^) and 8-*iso*-prostaglandin F_2α_ (8-*iso*-PGF_2α_) (*46*). Adropin overexpression using a lentivirus or pharmacological treatment with adropin peptide ameliorates palmitate-induced oxidative stress in hepatocytes through regulating Nrf2 and Nrf2-related antioxidative enzyme expression (*46*). Consistently, adropin therapy alleviates nonalcoholic steatohepatitis and liver injury progression through the CREB binding protein (CBP)-Nrf2-mediated antioxidative response pathway in mice (*45*). In line with these findings, Wu *et al*. demonstrated that adropin treatment reduces ischemia/reperfusion-induced myocardial injury in an H9c2 cardiomyoblast cell model and that these protective effects were through an increase of scavenging superoxide radical activity and inhibition of lipid peroxidation and inflammatory response (*25*).

In the present study, we found that post-ischemic treatment with adropin dramatically reduced gp91^phox^ (a significant source of O_2_^-.^generation) and the lipid peroxidation product, 4-HNE, in the ischemic cortex, both of which contribute to endothelial dysfunction since oxidative damage induced by gp91^phox^ or 4-HNE can impair eNOS function by oxidizing tetrahydrobiopterin (BH4), an essential co-factor for eNOS activity (*47*). In addition, reduced gp91^phox^ levels in adropin-treated animals potentially limits the reaction between NO and O_2_^-.^to form ONOO^-^, which in turn increases NO bioavailability in the ischemic brain. It is worth noting that ONOO^-^ is a highly reactive species known to activate matrix metalloproteinase (MMP)-9 (*41*) a protease involved in the degradation of extracellular matrix and tight junction proteins, thus leading to the BBB disruption (*20*). In support, we found that adropin treatment significantly reduced stroke-induced extravasation of IgG, albumin, and Hb (three BBB damage markers) along with degradation of tight junction proteins zona occludens (ZO)-1 and occludin (two major structural components of the BBB) in the ischemic cerebral cortex, and that these effects, mediated by adropin, were associated with a dramatic decrease of MMP-9 activity. Based on these results, we suggest that adropin-mediated neuroprotection in ischemic stroke might be mediated through activation of eNOS and reduction of oxidative stress, thus increasing NO bioavailability and inhibition of MMP-9 activity to maintain endothelial function and BBB integrity. Deficient production of eNOS-derived NO is the key factor contributing to endothelial dysfunction in diabetes, obesity, and hyperlipidemia (*48*). eNOS-derived NO decreases BBB permeability, reduces platelet aggregation, and limits leukocyte adhesion to the endothelium (*49-51*). We quantified levels of Ly-6B.2 (a specific neutrophil marker) in the ischemic brain since infiltrating neutrophils are the most important source of active MMP-9 in cerebral ischemia (*23*). Intriguingly, adropin treatment dramatically reduced Ly-6B.2 levels in the ischemic cortex induced by pMCAO. Thus, reduced MMP-9 activation by adropin treatment could result from reduced oxidative damage and/or reduced infiltration of active MMP-9-laden neutrophils. A schematic summarizing the proposed mechanisms through which adropin confers neuroprotection in stroke is presented in **Figure 8**.

**Figure 8.**
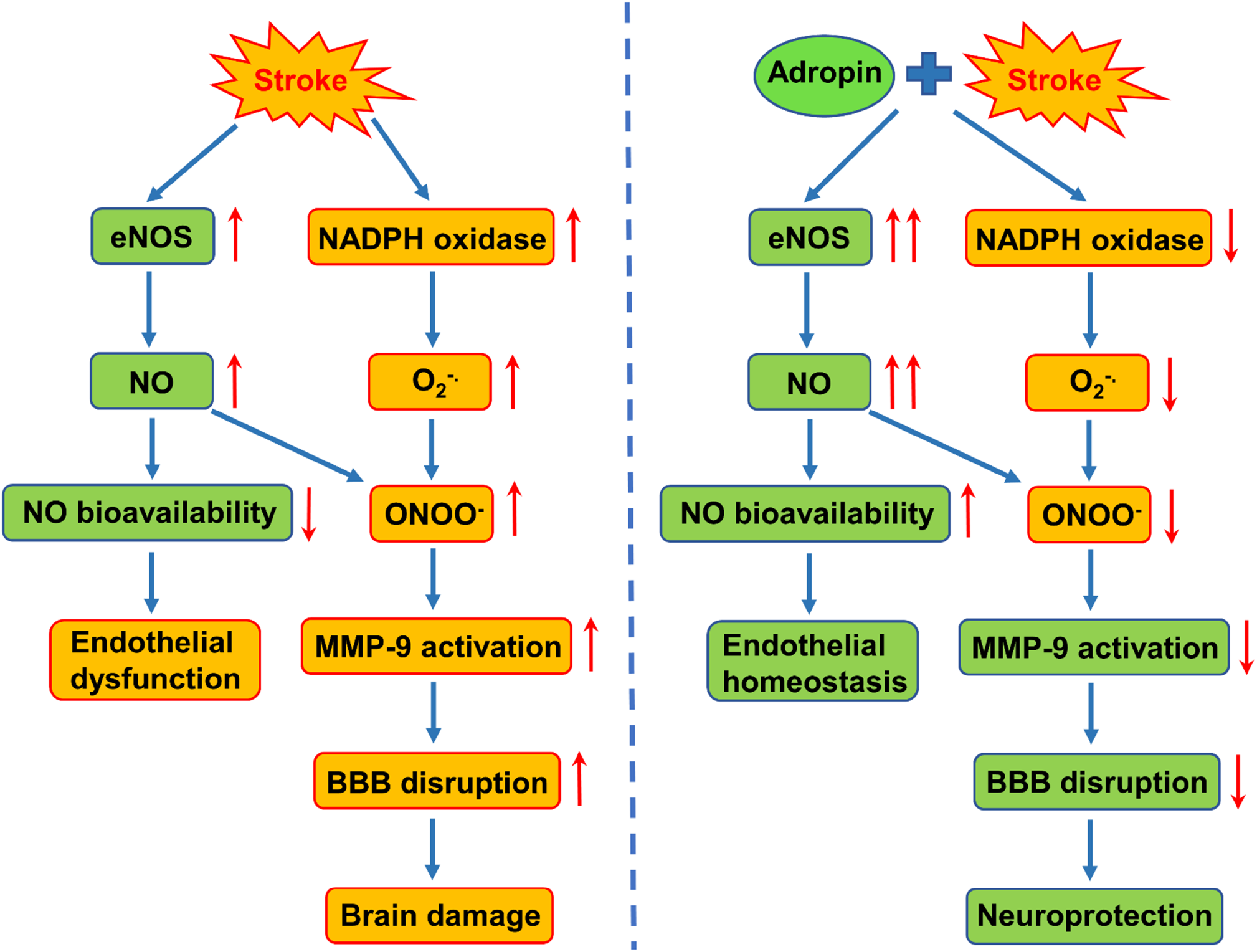
Proposed schematic mechanisms of adropin-mediated neuroprotection in ischemic stroke. ***Left*:** Stroke induces eNOS phosphorylation to release nitric oxide (NO), together with upregulation of NADPH oxidase (NOX) to produce superoxide (O2^-.^). The reaction between NO and O2^-.^ forms peroxynitrite (ONOO^-^), resulting in a decrease in NO bioavailability and MMP-9 activation through the S-nitrosylation of MMP-9 pro-domain “cysteine switch” mechanism, which ultimately leads to endothelial dysfunction, disruption of the blood-brain barrier (BBB), and brain damage. ***Right*:** Treatment with adropin or adropin overexpression can induce a higher level of eNOS phosphorylation-derived NO generation (double red arrow) and inhibit NOX from producing O2^-.^ in ischemic stroke, which decreases the ONOO^-^ formation. These effects may finally improve endothelial function and inhibit MMP-9 activity resulting in reduced BBB damage and neuroprotection.

### Limitations of the Study and Future Directions

There are several limitations of the current study. First, findings are restricted to permanent brain ischemia in mice. Considering that prompt reperfusion, either occurring spontaneously or following recanalization therapies, is only achieved in a minority of ischemic stroke patients (*52, 53*), in this study, we utilized a model of permanent ligation of the MCA resulting in cortical ischemia. Future studies are warranted to investigate the effects of adropin treatment on stroke outcomes in models of transient ischemia/reperfusion injury. We showed that adropin is protective in aged mice subjected to stroke. However, it will be important to expand these findings and evaluate the effects of adropin in stroke models with comorbidities. Diabetes is an independent risk factor for stroke (*54*) and hyperglycemia increases infarct volume, BBB disruption, and hemorrhagic transformation (*55*). Adropin plays a vital role in the modulation of insulin resistance, which potentially affects glucose metabolism (*8, 56*). Thus, it would be of great translational value to investigate the potential neuroprotective effects of adropin in animals with comorbid conditions such as diabetes or obesity subjected to ischemic stroke (transient and permanent MCAO models).

Another limitation is that the detailed molecular mechanisms underlying the biological effects of adropin are currently poorly understood as its receptor remains elusive. Although Stein *et al*. discovered GPR19 (an orphan G protein-coupled receptor) as a potential adropin receptor (*57*), a physical association between adropin and GPR19 itself has not been demonstrated. Another study showed that adropin interacts with neural recognition molecule-3 (NB-3)/contactin 6 (CNTN6), a brain-specific membrane-tethered Notch1 ligand (*9*). Both adropin receptor candidates, GPR19 and CNTN6, are highly expressed in the brain. More research is needed to better understand signaling mechanisms and establish the molecular identity of the adropin receptor, which would facilitate the therapeutic development of this peptide for the potential treatment of various diseases.

Brain adropin levels are significantly reduced in aged animals (*13, 14*), and following ischemic injury, as demonstrated in the current study. We previously showed a dramatic reduction in *Enho* transcription and adropin levels in brain endothelial cells exposed to hypoxia and low-glucose *in vitro* to mimic stroke conditions (*6*). Presently, the precise molecular mechanisms underlying the loss of brain adropin during the aging process and following ischemia remain undetermined. A possible mechanism is the proteolytic cleavage of the full-length adropin^1-76^ peptide by an unidentified protease/peptidase. Understanding how aging and ischemia alter adropin levels in the brain could shed new light on the biological function of this protein. It could also identify novel targets to prevent the loss of this neuroprotective peptide to enhance resilience to ischemic injury and other aging-associated brain diseases.

In summary, the present study fills an important gap in our mechanistic understanding of the effects of adropin in the context of ischemic brain injury. We show that post-ischemic treatment with exogenous adropin peptide or overexpression of endogenous adropin exerts beneficial effects on reduction of infarct volume and improvement of long-term neurological deficits in both young and aged mice subjected to ischemic stroke and that these neuroprotective effects of adropin are most likely mediated through activation of an eNOS-dependent pathway.

## Methods

All animal procedures complied with the National Institutes of Health (NIH) guidelines and were approved by the University of Florida Institutional Animal Care and Use Committee under animal protocol number 201907934.

### Animals

Mice were bred and housed in a temperature- and humidity-controlled animal facility with a 12-hour light/dark cycle. Food and water were available *ad libitum*. Male and female adropin transgenic (AdrTg) hemizygous mice (*8*) and heterozygous carriers of the null adropin allele (*Enho*^*+/-*^) (*18*) on the C57BL/6J background were maintained in an AAALAC-accredited, germ-free facility at the University of Florida. Wild-type (WT) littermates from both strains were used as controls. Male 10- to 12-week-old C57BL/6 mice were purchased from Taconic Biosciences (Hudson, NY) and eNOS knockout (*eNOS*^*-/-*^) mice (Stock No: 002684) were purchased from The Jackson Laboratory (Bar Harbor, ME, USA). Mice were assigned to experimental groups using the GraphPad randomization tool and coded at the time of surgery to guarantee a blinded analysis of the data. A total of 510 mice were utilized in the current study.

### Genotyping

Genomic DNA was extracted from tail tissues and amplified with specific PCR primers using REDExtract-N-Amp™ Tissue PCR kit (Cat. No. XNAT-100RXN, Sigma-Aldrich, Saint Louis, MO) according to the manufacturer’s instructions. The PCR reaction was run under the following conditions: 94°C for 3 min, followed by 35 cycles at 94°C for 30s, 60°C for 30s, and 72°C for 45s, followed by a 10-min 72°C final extension. PCR products were electrophoretically separated on a 2% agarose gel. As a result, genotyping of AdrTg mice was performed on tail tissue genomic DNA by PCR using one pair of primers (forward: 5’-GGCTATTCTCGCAGGATCAGT-3’; reverse: 5’-GAGAGCCCCTTGGGAGATG-3’) from the human β-actin promoter and *Enho* gene coding region, where the 80 bp amplicon confirms the presence of the transgene (overexpression of adropin). Genotyping of wild-type (*Enho*^*+/+*^, 72 bp), heterozygous (*Enho*^*+/-*^, 72 and 170 bp), and homozygous (*Enho*^*-/-*^, 170 bp) was performed using genomic DNA obtained from the mouse tail by PCR using two pairs of *Enho* wild-type (forward: 5’-GGAAGATGTACGTTCAGACACACA-3’; reverse: 5’-GTGTGCCTAGGTGTACCTTTCC-3’) and excised (forward: 5’-ACGTTCAGACACACACCTGATC-3’; reverse: 5’-CTACAATCTAGGCTGCTGGGT-3’) primers.

### Surgical procedures and treatments

Permanent focal cerebral ischemia was induced by ligation of the left middle cerebral artery (MCA), as described previously (*58*). Briefly, mice were anesthetized with isoflurane in medical grade oxygen (3% for induction and 1.5-2% for anesthesia maintenance during surgery). Mice were given 0.05 mg/kg of buprenorphine hydrochloride subcutaneously pre-surgery and every 12h for the first two days after stroke to minimize pain and distress. Once the surgical level of anesthesia was reached, the left common carotid artery (CCA) was exposed and carefully dissected from the vagus nerve. Then a 6-0 silk suture was applied to ligate the left CCA permanently. Subsequently, a skin incision was made between the eye and the left ear under a stereomicroscope. The temporal muscle was retracted to locate the left MCA via skull transparency. A small round craniotomy (∼1-1.5mm in diameter) was carefully performed between the zygomatic arch and the squamosal bone to expose the MCA using a surgical drill burr (Cat. No. 726066) connected to a battery-operated Ideal Micro-Drill (Cat. No. 726065; Cellpoint Scientific, Gaithersburg, MA). Finally, two to three drops of sterile saline were applied to the target area, and the meninges covering the MCA were carefully removed with a fine forceps. Before the bifurcation between the frontal and parietal MCA branches, the MCA distal trunk was permanently ligated using an ophthalmic 9-0 silk suture. A clear interruption of blood flow was visually confirmed. After surgery, mice were allowed to recover in a temperature-controlled chamber. Sham-operated mice received the same surgical procedures except for the MCA occlusion. In the experiments involving exogenous adropin peptide treatment, mice were randomized to receive either saline or a single injection of synthetic adropin peptide (90-2700 nmol/kg) at the onset of ischemia or 3h after pMCAO via the external jugular vein using a 31-gauge needle.

### Inclusion and exclusion criteria

Animals were allocated to experimental groups and included in the final statistical analysis following predefined inclusion and exclusion criteria. A successful MCA occlusion was confirmed during surgery by visual inspection of the artery showing successful interruption of the blood flow for at least one minute. Animals were excluded from the study if the brain’s surface was physically damaged by forceps or the microdrill. Hemorrhage, more than 20% body weight loss, or death before the endpoint were additional exclusion criteria. In our hands, the pMCAO model of ischemic stroke results in extremely low mortality. In the current study, three adult wild-type mice and two aged AdrTg mice died before the end of the study.

### Measurement of infarct volume

The infarct volume was assessed by either 2,3,5-triphenyltetrazolium <u>c</u>hloride (TTC), MRI, or Cresyl violet staining at 48 hours, 14 days, or 21 days after ischemic stroke, respectively.

#### TTC staining

Forty-eight hours after pMCAO, mice were euthanized and perfused transcardially with ice-cold saline, and brains were harvested. Brains were sliced into six 1-mm-thick coronal sections. The fourth section starting from the rostral side, was dissected into ipsilateral and contralateral cerebral cortices. Tissues were immediately frozen in liquid nitrogen and saved in the −80°C freezer until later processing for molecular biology analyses. The remaining sections were stained with 2% TTC in phosphate-buffered saline (PBS, pH 7.4) at room temperature for 30 min, followed by fixation with 4% paraformaldehyde (PFA) in PBS, pH 7.4. Sections were scanned at 600 dpi rostral side down using an HP Scanjet 8300 scanner (Palo Alto, CA) and saved as a JPEG file. Also, the caudal side of the third section, which corresponds to the rostral side of the fourth coronal section, was scanned to measure the infarct area of the fourth slice. Infarct volume was determined using Adobe Photoshop CS5 as described in detail in our previous study (*58*).

#### Cresyl violet staining

Briefly, mice were euthanized at 21 days after pMCAO and transcardially perfused with 10 mL of saline containing 5 mM EDTA using a peristaltic pump at a speed of 5 mL/min, followed by 30 mL of 4% PFA in PBS. Brains were removed and post-fixed in 4% PFA for 24 hours, then transferred to PBS for storage at 4°C. Brains were cut into a series of 30-µm thick coronal sections on a semi-automated vibrating microtome (Compresstome^®^ Model No. VF310-0Z; Precisionary Instruments, Natick, MA). Cortical infarct volume was quantified from infarcted areas in ten brain sections spaced 0.5 mm apart using Cresyl violet staining, as described in previous studies (*59*). Sections were scanned using an Aperio ScanScope^®^ CS system and analyzed with ImageScope Software (Aperio Technologies, Vista, CA). The border between infarcted and non-infarcted areas was outlined.

#### Magnetic Resonance Imaging (MRI)

Mice were euthanized at 14 days after pMCAO and perfused transcardially with saline followed by 4% PFA in PBS. Heads were removed and post-fixed in 4% PFA for 24 hours, then transferred to PBS for storage at 4°C until imaging by MRI. Mouse heads were brought to room temperature overnight and moved to a 15 mL conical tube containing FC-40 oil. Heads were aligned with snouts parallel to the tip of the conical tube and the caudal occipital region secured with a custom-fitted plastic holder. The entire setup was sealed with paraffin, and care was taken to minimize the presence of visible air pockets. Images of fixed excised mouse heads were collected in an 11.1 Tesla MRI scanner (Magnex Scientific Ltd., Oxford, UK) with a Resonance Research Inc. gradient set (RRI BFG-240/120-S6, maximum gradient strength of 1000 mT/m at 325 Amps and a 200 µs risetime; RRI, Billerica, MA) and controlled by a Bruker Paravision 6.01 console (Bruker BioSpin, Billerica, MA). An in-house 2.0 cm x 3.5 cm quadrature radiofrequency (RF) surface transmit/receive coil tuned to 470.7 MHz (^1^H resonance) was used for B1 excitation and signal detection (RF engineering lab, Advanced Magnetic Resonance Imaging and Spectroscopy Facility, Gainesville, FL). A T2-weighted Turbo Rapid Acquisition with Refocused Echoes (TurboRARE) sequence was acquired with the following parameters: effective echo time (TE)= 20 ms, TE=5 ms, repetition time (TR)= 4 seconds, RARE factor= 8, number of averages= 6, field of view (FOV) of 15 mm x 15 mm and 0.5 mm thick slice, and a data matrix of 256 × 256 and 26 interleaved ascending coronal (axial) slices (left-right orientation) covering the entire brain from the rostral-most extent of the prefrontal cortical region and caudally to the upper brainstem/cerebellum. Saturation bands were applied around the brain to suppress non-brain (muscle) signal. Whole-brain and lesion boundary masks were generated using image contrast and manual segmentation tools available in ITKSNAP (*60*). Lesion volumes were exported for statistical analyses. Representative brain scans were further processed for 3-dimensional visualization in BrainNet viewer in MATLAB (*61*). Brain extraction was carried out using image math programs in FMRIB Software Library (FSL 6.0.3) (*62*). N4 bias field correction (*63*) was used to remove B1 RF field inhomogeneities and reduce FOV intensity variations in T2 anatomical scans before alignment to the mouse brain template (common coordinate framework, version 3) (*64*). Cropped T2 TurboRARE scans were first linearly registered to the mouse brain template using FSL FLIRT, and then in ANTs, we used antsRegistrationSynQuick.sh to warp images to the same template. Linear and warping matrices were stored and applied to the lesion volume masks.

### Neurobehavioral tests

A battery of neurobehavioral tests, including open field test, cylinder test, corner test, nesting test, adhesive removal test, and novel object recognition, were performed before (baseline) and at several defined time points after stroke to assess sensorimotor and cognitive deficits of ischemic stroke mice.

#### Open field test

The open field test assesses motor function and exploratory locomotion (*58*). Mice were individually placed in an open field chamber (40 × 40 × 40 cm) with grey sidewalls and allowed to explore it for 10 min. After each test, the open field arena was cleaned with 70% ethanol. Movements of animals were recorded with a video camera placed above the apparatus and analyzed using Any-maze software (Stoelting, Wood Dale, IL). The total distance traveled in the arena was analyzed.

#### Cylinder test

The cylinder test was used to assess forepaw use asymmetry (*65*). Mice were placed in a transparent cylinder (diameter: 9.5 cm, height: 30 cm; Cat. No. B0070ZROK) supported by a piece of transparent glass on a larger cylinder. A video camera placed in the larger cylinder below recorded all forelimb movements of the mouse in the top cylinder for 5 min. At least 10 instances of complete rearing and 20 times of forelimb contacts are needed to reach the criterion for analyzing the use of forelimbs (left/right/both) in slow motion by the VideoLAN Client (VLC) media player software (version 3.0.11). The first contact events (%, with both forelimbs) were analyzed when the first contact on the cylinder wall happened with both forelimbs during the particular full rearing period. The contact with both forelimbs is when the mouse contacts the cylinder wall simultaneously (the interval between both forelimbs contacts is no more than 3 frames at 29 frames per second). The calculation for the first contact events with both forelimbs (%) = total number of first contact events with both forelimbs/total number of full rearings with any forelimb contact (s) × 100.

#### Corner test

The corner test was performed to assess overall sensorimotor and postural asymmetries as described in a previous study (*59*). The injured mouse turns preferentially toward the contralateral (right). Each mouse performed a total of 12 trials with at least 5 seconds intervals between each trial. Performance was expressed as the number of left turns out of 12 trials for each test.

#### Nesting test

Nest building is an innate behavior that assesses sensorimotor function in mice (*66*). Briefly, one cotton nestlet (approximately 2 g) was placed into the cage 1h before the start of the dark cycle. On the morning of the next day, the nest was captured for scoring, and the weight of the unshredded nestlet was determined. The nest quality was assessed on a 0-5 rating scale, with 0 meaning the nestlet was not touched at all; 1, nestlet more than 90% intact; 2, nestlet was partially torn but 50-90% intact; 3, nestlet was torn with less than 50% remaining intact, but no obvious nest with shreds spread around the cage; 4, a flat nest without raised walls, more than 90% of the nestlet was shredded; 5, the nest was built perfectly with walls higher than mouse height and no unshredded pieces.

#### Adhesive removal test

This test assesses sensorimotor function, including forepaw and mouth sensitivity (time-to-contact) and dexterity (time-to-remove) as described previously by our group and others (*22, 67*). Briefly, mice were placed in a clean empty cage for at least 1 min to habituate to the testing condition, then the test mouse was restrained, and forceps were carefully used to place a white, round sticker tape (3/16 inches ≈ 5mm in diameter; Cat. No. 9787-8690 from USA Scientific Inc.) to the ventral surface of the contralateral forepaw. Subsequently, mice were quickly released back to the cage. First, the experimenter recorded the latency to touch the sticker (time-to-contact) and then remove the sticker (time-to-remove) by a mouse with its mouth up to a cut-off time of 2 min. Animals were pre-trained one day before surgery, and 5 trials were tested on each mouse after they learned how to remove the sticky tape using their mouths. The average time of 5 trials was used as the pre-surgery baseline value. At defined time points after stroke, mice were tested for another 5 trials, and the mean latency was used for analysis.

#### Novel object recognition

The novel object recognition (NOR) test evaluates long-term recognition memory (*68*). Briefly, for the training session, mice were individually placed in an open field chamber (40 × 40 × 40 cm) with grey sidewalls containing two identical objects for 8 minutes and returned to their home cage. After 24 hours, the mice were exposed to the familiar arena in the presence of the original object and a novel object to test long-term recognition memory for another 8 min. During the 8 min test session, the time spent exploring each object was recorded. The discrimination of recognition novelty was determined as a discrimination index: (time exploring the new object − time exploring the old object)/ (total time exploring both objects). Movements of animals were videotaped by an overhead camera and analyzed using the Any-maze software.

### Protein extraction and Immunoblotting

At the time of euthanasia, mouse brains were harvested following transcardiac perfusion with ice-cold saline, and cortical tissues were collected quickly, immediately frozen in liquid nitrogen, and stored in the −80°C freezer until further processing. Cortical tissues were homogenized in modified radioimmunoprecipitation (RIPA) lysis buffer consisting of 50 mM Tris-HCl (pH 7.4), 150 mM NaCl, 5 mM EDTA, 1 mM EGTA, 1% NP-40, 0.5% sodium deoxycholate and 0.1% SDS plus protease and phosphatase inhibitor cocktails (Cat. Nos. 78430 and 78428, respectively; Thermo Fisher Scientific, Rockford, IL). Tissue homogenates were sonicated and centrifuged as described in detail in our previous reports (*13, 69*). Protein concentration in the resulting tissue lysate supernatants was measured using the Pierce™ BCA assay kit (Cat. No. 23227, Thermo Scientific, Rockford, IL), and samples were aliquoted and stored at −80°C until analysis.

Fifty micrograms of protein lysates were denatured in 2× Laemmli’s sample buffer containing 4% β-mercaptoethanol at 70°C for 10 min for measuring most of the phosphorylated and total protein levels, while non-reducing conditions were used to measure proteins related to oxidative stress including gp91^phox^ (the catalytic subunit of NADPH oxidase) and 4-hydroxynonenal (4-HNE) (*13*), and a native condition (non-reduced and non-denatured) was used to detect a specific neutrophil marker Ly-6B.2 (*58*), as we described previously. All samples were separated on 4-20% SDS-PAGE and then transferred onto nitrocellulose membranes. Membranes were then blocked for 1h at room temperature with 5% non-fat milk (for eNOS, ZO-1, Occludin, MMP-9, gp91^phox^, 4-HNE, CD31, Akt, ERK1/2, and β-actin) in Tris-buffered saline or Odyssey blocking buffer (Cat. No. 927-50000; Li-Cor, Lincoln, NE; for adropin, peNOS^Ser1176^, pAkt^Ser473^, pERK1/2, Claudin-5, and Ly-6B.2). After blocking, membranes were incubated overnight at 4°C with antibodies against either adropin, peNOS^Ser1176^, pAkt^Ser473^, pERK1/2, eNOS, Akt, ERK1/2, CD31, ZO-1, Occludin, Claudin-5, MMP-9, gp91^phox^, 4-HNE, Ly-6B.2, or β-actin. After incubation with primary antibodies, membranes were washed with TBST 3 times and then incubated for 1h with goat anti-rabbit IRDye 800CW, goat anti-mouse IRDye 800CW, or donkey anti-mouse IRDye 680LT secondary antibodies. Membranes were scanned with an Odyssey infrared scanning system (Li-Cor), and immunoreactive bands were quantified using the Image Studio software (Li-Cor). The vendor, catalog number, and working dilution of all the antibodies utilized in this study are included in **Supplemental Table 2**.

### Immunohistochemistry

Mice were transcardially perfused with saline containing 5 mM EDTA followed by 4% PFA in PBS. Brains were harvested and stored in 4% PFA at 4°C. After 24 hours of incubation, brains were transferred to PBS and kept at 4°C for less than one week until sectioned at 20 µm using a semi-automated vibrating microtome. Double immunostaining of adropin with different cell markers was performed following our previously described protocol (*70*). Briefly, brain sections were mounted to a slide and dried at 50°C for 30 min to ensure that tissue adhered tightly to the slide. Mounted tissue was permeabilized with tris-buffered saline (TBS) with 0.1% Triton X-100 for 5 min. The sections were subsequently blocked for 1h at room temperature in TBS solution with 0.05% Triton X-100 (TBST) containing 1% bovine serum albumin and 10% normal goat serum, followed by overnight incubation at 4°C with the following primary antibodies prepared in antibody diluent (TBS solution with 0.02% Triton X-100 plus 1% bovine serum albumin and 2% normal goat serum): mouse anti-adropin (Cat. No. 14117, 1:200; Cayman Chemical), rat anti-CD31 (Cat. No. NB600-1475, 1:100; Novus Biologicals, LLC, Littleton, CO), rabbit anti-NeuN (Cat. No. NBP1-77686, 1:100; Novus Biologicals), rabbit anti-GFAP (Cat. No. Z0334, 1:100; Dako North America, Inc., Carpinteria, CA), rabbit anti-Iba1 (Cat. No. 019-19741, 1:100; Wako Chemicals USA, Inc., Richmond, VA). After washing with TBST, sections were incubated for 90 min at room temperature with goat anti-mouse Alexa Fluor 594 (Cat. No. A11032, 1:250; ThermoFisher Scientific, Waltham, MA) and either of goat anti-rat Alexa Fluor 488 (Cat. No. A11006, 1:250; ThermoFisher Scientific) or goat anti-rabbit Alexa Fluor 488 (Cat. No. 111-545-144, 1:250; Jackson ImmunoResearch Laboratories, West Grove, PA). Alternate sections from each experimental condition were incubated in all solutions except the primary antibodies to assess nonspecific staining. Sections were then counterstained with 100 nM 4′,6-diamidino-2-phenylindole (DAPI) in TBS for 10 seconds at room temperature and coverslipped with Fluoromount™ aqueous mounting medium (Cat. No. F4680; Sigma-Aldrich). Fluorescence images were captured with a spinning disk confocal microscope (Cat. No. DSU-IX81; Olympus, Center Valley, PA).

### RNA extraction and quantitative real-time PCR

Total RNA from the mouse brain cortex was isolated using a modified method of acid guanidinium thiocyanate-phenol-chloroform extraction (*71*). RNA concentration and purity were measured by absorbance using a Take3 Micro-Volume Plate Reader (Biotek Instruments, Winooski, VT). According to the manufacturer’s instructions, total RNA (1 µg) was transcribed into cDNA using ProtoScript® II First Strand cDNA Synthesis Kit (Cat. No. E6560; New England BioLabs). Quantitative real-time PCR was performed with 20 ng of cDNA in a total reaction volume of 10 μL using Luna® Universal qPCR Master Mix (Cat. No. M3003L; New England BioLabs) according to the manufacturer’s protocol. The primer sequences used for amplification of mouse *Enho* gene were: forward, 5′-ATGGCCTCGTAGGCTTCTTG-3′; and reverse, 5′-GGCAGGCCCAGCAGAGA-3′ and were normalized to the housekeeping gene *Hprt1*: forward, 5′-CCCCAAAATGGTTAAGGTTGC-3′; and reverse, 5′-AACAAAGTCTGGCCTGTATCC-3′. PCR reactions were run in triplicate, and cycle threshold values were normalized to *Hprt1* expression for each sample.

### Measurement of adropin level in plasma

To determine the stability of synthetic adropin peptide following intravenous injection, 900 nmol/kg exogenous adropin peptide was administered to mice via the jugular vein. 20-30 µL of blood were collected in EDTA-coated tubes (Cat. No. NC9414041, Fisher Scientific) before injection (basal level) and at 5 min, 1 h, 4 h, and 24 h after injection. Blood was immediately spun down at 2,000×g for 10 min to save the plasma for adropin ELISA assay. Adropin levels in mouse plasma were quantified using an ELISA kit (Cat. No. EK-032-35, Phoenix Pharmaceuticals, Inc., Burlingame, CA) as recommended by the manufacturer’s protocol. Plasma collected at 5 min after injection was diluted 5000 times with assay buffer (since synthetic adropin peptide in plasma is highly concentrated at this time point). Plasma collected before injection (baseline) and at 1 h, 4 h, and 24 h after injection was diluted 50 times. A total volume of 50 µl of diluted samples or standard synthetic adropin peptide (0.01-100 ng/mL) was added to the appropriate microtiter wells. For plasma collected after 48 hours from stroke mice treated with 0.1% BSA in saline or 900 nmol/kg adropin synthetic peptide, plasma was diluted 10 times with assay buffer, and 50 µL of diluted samples was used for the ELISA assay. All standards and samples were assayed in duplicate, and optical absorbance at 450 nm was measured with a Synergy™ HT Multi-Mode Plate Reader (Biotek Instruments, Winooski, VT). The concentrations of adropin in plasma samples were determined from a four-parameter logistic curve fitted for the standard peptide absorbance values.

### Measurement of nitrite and nitrate levels in plasma

Due to nitric oxide’s instability *in vivo*, nitrite and nitrate levels serve as surrogates of NO production. Levels of nitrite/nitrate in plasma were measured using a commercially available kit (Cat. No. 780001, Cayman Chemical, Ann Arbor, MI) as instructed in the manufacturer’s manual. Thirty minutes after treatment with either adropin synthetic peptide (900 nmol/kg; i.v.) or vehicle (0.1% BSA in saline), blood was collected in EDTA-coated tubes (Cat. No. NC9414041, Fisher Scientific) and immediately spun down at 2,000×g for 10 min. Then plasma was saved in liquid nitrogen and kept at −80°C until use. 80 µL of thawed plasma was mixed with an equal volume of assay buffer provided with the kit, then filtered through a 3 kDa molecular weight cut-off filter (Cat. No. OD003C33, Pall Corporation, Show Low, AZ) at 14,000 ×g for 15 min and the filtrates were saved. Finally, 80 µL of the filtrates or nitrite standards were loaded to a 96-well plate, mixed with the Griess reagent, and absorbance was measured at 540 nm on a Synergy™ HT Multi-Mode Plate Reader. The concentrations of nitrite/nitrate in plasma samples were calculated from a standard curve prepared with nitrite standards.

### MMP-9 activity assay

MMP-9 enzymatic activity in the cortical homogenate was measured using our group’s fluorescence resonance energy transfer (FRET) peptide immunoassay (*72*). Briefly, 96-well high binding plates (Cat. No. 655077; Greiner Bio-One, Monroe, NC) were coated with MMP-9 antibody (Cat. No. sc-6841R) following protein A/G pre-coating. Then, a total of 50 μg protein extracted from the mouse brain cerebral cortex was added to each well and incubated at 4°C overnight. After incubation, wells were washed with TCNB buffer (50 mM Tris, 10 mM CaCl_2_, 150 mM NaCl, 0.05% Brij-35) and 1 µM of 520 MMP FRET substrate III (Cat. No. 60570-01; AnaSpec, San Jose, CA) was added. Plates were incubated for 48h at 37°C, then relative fluorescence units (RFUs) were read at excitation/emission wavelengths of 485/528 nm in a Synergy™ HT Multi-Mode Plate Fluorescence Reader. The average value from one paired substrate control wells was used to subtract baseline fluorescence from sample wells.

### Assessment of blood-brain barrier permeability by ELISAs

Brain levels of immunoglobulin G (IgG), albumin and hemoglobin (Hb), three sensitive BBB damage markers, were measured using commercially available ELISA kits (Cat. Nos. E90-131 and E99-134 for IgG and Albumin, Bethyl Laboratories, Inc., Montgomery, TX; Cat. No. E-90HM for Hb, ICL, Inc., Portland, OR) according to the manufacturer’s instructions. A total of 50 µg, 3 µg, and 10 µg of protein extracted from the ipsilateral and contralateral cerebral cortex of the mouse brain were used for the IgG, Albumin, and Hb measurement, respectively. All samples were assayed in duplicate, and optical absorbance was measured at 450 nm with a Synergy™ HT Multi-Mode Plate Reader.

### Vessel casting, visualization of cerebrovasculature, and analysis of anastomosis

Cerebrovascular anatomy was quantitatively evaluated in AdrTg, *Enho*^*-/-*^ mice, and their wild-type littermates to assess the potential impact of variations in the anatomy of the cerebral circulation among mouse strains on susceptibility to ischemic injury. Mice were deeply anesthetized with 2.5-3% isoflurane, and 50 mg/kg papaverine hydrochloride dissolved in saline was administered via an external jugular vein. After 20-30 seconds to allow the vessels to be fully dilated, the heart was quickly exposed, and the descending aorta was clamped using hemostatic forceps. The right atrium was opened, and 1 mL of 1000 U/mL heparin was injected transcardially, followed by 3 mL of blue latex rubber solution (Cat. No. BR80B; Connecticut Valley Biological Supply Co., Southampton, MA) with a flow rate of ∼1 mL/min. After that, the brain was gently taken out from the skull and fixed in 4% PFA for 24 hours at 4°C before imaging the dorsal and ventral surface with an HP Scanjet 8300 scanner at 2400 dpi. Anastomoses on the dorsal surface of the hemispheres were localized by tracing the distal branches of the anterior cerebral artery (ACA) and the MCA to the anastomosis points (*73*). Adjacent anastomosis points were connected by the line of anastomoses. The distance from the midline to the line of anastomoses was measured in coronal planes 2, 4, and 6 mm from the frontal pole in photographs taken from the dorsal brain surface using Adobe Photoshop CS5 software.

### Two-dimensional laser speckle imaging

Cortical cerebral blood flow (CBF) was monitored using a two-dimensional laser speckle contrast image system (MoorFLPI-2; Moor Instruments Inc. Delaware) as previously described (*74*). Briefly, anesthetized mice were placed in the prone position with the skull exposed but unopened. CBF was recorded in both cerebral hemispheres immediately. After 10 min of recording, saline, acetylcholine (ACh, 1.0 mM solution, 100mL/10g of body weight), or synthetic adropin peptide (900 nmol/kg) was administered intravenously from the tail vein, and the CBF was recorded continuously for another 30 min. Representative CBF images at the selected time-points from the entire recording period include the arbitrary units in a 16-color palette calculated by the MoorFLPI software. The average CBF change (CBF %) within the total 30-min recording was normalized to the baseline CBF (first 10 min).

### Isolation of brain microvessels

Brain microvessels were isolated from individual mouse brains as described in a recent report with modifications (*75*). After removal of the brain stem and cerebellum, brain hemispheres were transferred to a 2 mL reinforced polypropylene tube (Cat. No. 330TX; BioSpec Products, Inc., Bartlesville, OK) pre-filled with 1 mL of homogenization buffer (101 mM NaCl, 4.6 mM KCl, 2.5 mM CaCl_2_, 1.2 mM KH_2_PO_4_, 1.2 mM MgSO_4_, 15 mM HEPES, pH 7.4) with phosphatase/protease inhibitor cocktails and immediately frozen in liquid nitrogen, and kept in the −80°C freezer until further processing. After thawing, 1.8 g of stainless beads (3.2 mm, Cat. No. 11079132ss; BioSpec Products, Inc.) were carefully added to the reinforced tube, and the brain tissue was homogenized by a bead homogenizer (Bead Mill 4, Thermo Fisher Scientific) for 60 s at a speed of 3 m/s. Then, the homogenate was transferred to a new 2 mL Eppendorf tube and centrifuged (1000×g, 10 min, 4°C). Before centrifugation, a part of the brain homogenate (50 µL) was collected in another tube as a whole-brain fraction. After centrifugation, the supernatant was removed carefully, and 1.5 mL of 18% (w/v) dextran (molecular weight: 75,000 kDa; Cat. No. DE125; Spectrum Chemical) was added to the pellet and mixed well using a pipette. The samples were immediately centrifuged (10,000×g, 15 min, 4°C), and the suspended myelin and supernatant were carefully discarded. Pellets were resuspended in 400 µL of suspension buffer (homogenization buffer containing 25 mM NaHCO_3_, 10 mM glucose, 1 mM pyruvate, and 5 g/L bovine serum albumin) and dispensed with a pipette. The samples were filtered through a cell strainer (70 µm) (Cat. No. 43-10070-50; pluriStrainer-Mini, pluriSelect, Leipzig, Germany). The strainer mesh was washed 6 times with 400 µL of suspension buffer. Samples passed through the 70 µm mesh were added to a cell strainer filled with 800 mg of glass beads (0.5 mm, Cat. No. 11079105; BioSpec Products, Inc.) placed on 15 mL Falcon tube and then washed 10 times with 500 µL of suspension buffer. After washing, glass beads were transferred to a new 2 mL Eppendorf tube using a plastic spatula, and 1 mL of suspension buffer was added and mixed by inverting. The supernatant was then quickly transferred into a new 2 mL Eppendorf tube. Glass beads were re-added into the 1 mL of suspension buffer and mixed by inverting. The supernatant was transferred rapidly into the previous tube, and part of the brain capillary fraction was used for microscopy. The tube was centrifuged (10,000 ×g, 10 min, 4°C), and the supernatant was removed. The pellet was resuspended in 100 µL of RIPA buffer containing phosphatase/protease inhibitor cocktails by sonication to extract protein as described in detail in the section “Protein extraction and Immunoblotting”. Protein concentration was measured using a Pierce BCA protein assay kit. The enrichment of the brain microvessels compared to total brain homogenate was analyzed by western blotting with antibodies against endothelial markers such as CD31, eNOS, and tight junction proteins such as ZO-1 and claudin-5.

### Power analysis, sample size calculation, blinding, and randomization

Using the G*Power v.3.1 software, we performed an *a priori* sample size calculation for each outcome measurement. To estimate the required sample size, we utilized variances from our preliminary studies in the pMCAO mouse model to calculate effect size comparing two independent groups using α=0.05 and β (type II error) of 0.2 with a power of 80%. This study was powered to detect at least a 25% difference between vehicle- and adropin-treated groups, which is a biologically meaningful effect (*76*). As calculated, no less than 6 mice per group were needed for the measurement of infarct volume and neurobehavioral deficits, and 3-5 mice per treatment group or genotype were required for molecular studies; 5-7 mice per group with different genotype background were required for physiological parameters measurement. All mice were coded and randomly allocated to experimental groups using the randomization tool developed by GraphPad Prism (http://www.graphpad.com/quickcalcs/randomize1.cfm). The investigators performing stroke surgeries, euthanizing animals, or performing outcome assessments (behavioral tests, infarct size calculation, or molecular biology analyses) had no knowledge of the treatment group or genotypes to which an animal belonged.

### Statistical analysis

Data were analyzed using GraphPad Prism version 6 (GraphPad Software, San Diego, CA). An independent unpaired Student’s t-test was performed for comparison of two groups. Multiple comparisons were made using one-way, or two-way ANOVA followed by Bonferroni’s post hoc test. Values were expressed as mean ± SEM, and a *p-*value of less than 0.05 was considered statistically significant.

## Acknowledgments

Authors are grateful for the support from the AMRIS facility and the microscopy core at the McKnight Brain Institute, University of Florida.

## Author contributions

Conceptualization: CY, AAB, ECJ

Methodology: CY, BL, LL, BDS, KMD, JL, MP, MF, YYS, YMK, ECJ

Software: BDS, MP, MF

Formal analysis: CY, LL, ECJ

Data curation: CY, ECJ

Visualization: CY, AAB, ECJ

Writing – original draft: CY, MF, ECJ

Writing – review and editing: CY, JL, AAB, MM, ECJ

Supervision: MF, MM, CYK, SAF, AAB, ECJ

Funding acquisition: CY, AAB, ECJ.

## Data and materials availability

All data needed to evaluate the conclusions in the paper are available in the main text and the Supplementary Materials.

