## Supplemental Data for "Adropin confers neuroprotection and promotes functional recovery from ischemic stroke"

**This PDF file includes:**

Figs. S1 to S7

Tables S1 to S3

Full unedited Western blots

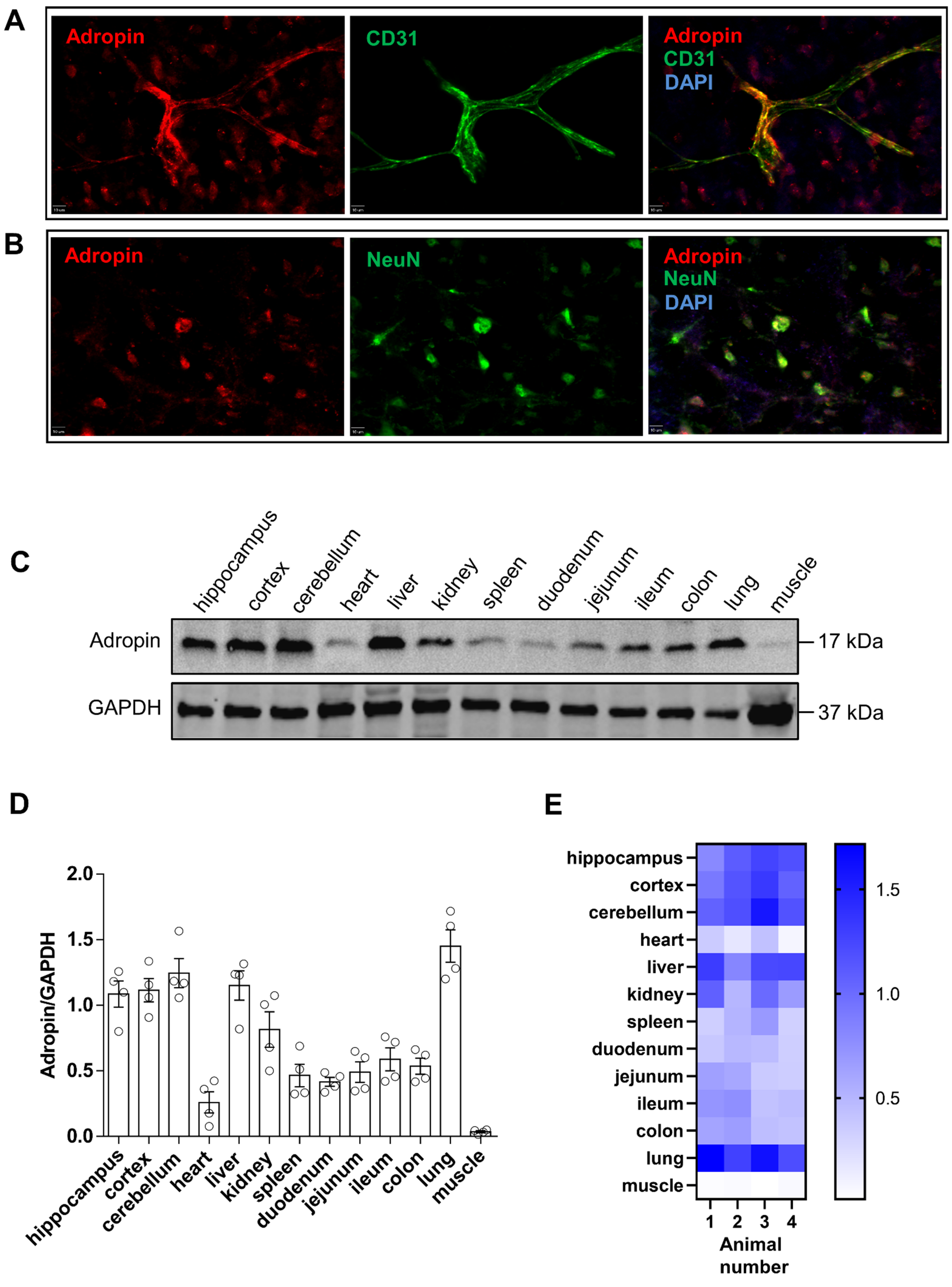


**Supplemental Figure 1. Immunohistochemical analysis of adropin expression in the mouse brain and levels of adropin in peripheral organs. A, B**, Representative confocal microscopy for adropin (red), endothelial cells (CD31, green), and neurons (NeuN, green) as well as nuclei (DAPI, blue) in naïve mouse brain. Adropin and CD31 double labeling show that adropin is highly expressed on endothelial cells in the cerebral cortex (**A**). Also, adropin is present on NeuN immunoreactive cells in the cortex (**B**). Scale bars: 10 $\mu$m. Data are representative of three independent experiments. **C,** Representative western blot for adropin in homogenates from different organs of adult (10-12 weeks) naïve male C57/BL6J mice with GAPDH as a loading control. **D, E,** Densitometric analysis (**D**), and heatmap (**E**) show that adropin is highly expressed in endothelial vessel-enriched organs including brain, liver, kidney, and lung, moderately expressed in spleen and intestines, and expressed at very low levels in the heart and skeletal muscle. n=4 per group.

A

| | |
|---|---|
| **Sequence:** | [NH2]CHSRSADVDSLSESSPNSSPGPCPEKAPPPQKPSHEGSYLLQP[COOH] |
| **Modification:** | [cyclo(S-S)] |
| **Project Number:** | CMB102728.1 |
| **Reference Number:** | A1314-1 |
| **Peptide Name:** | adropin |
| **Molecular Weight:** | 4499.90 |
| **Amount:** | 100 mg |

B

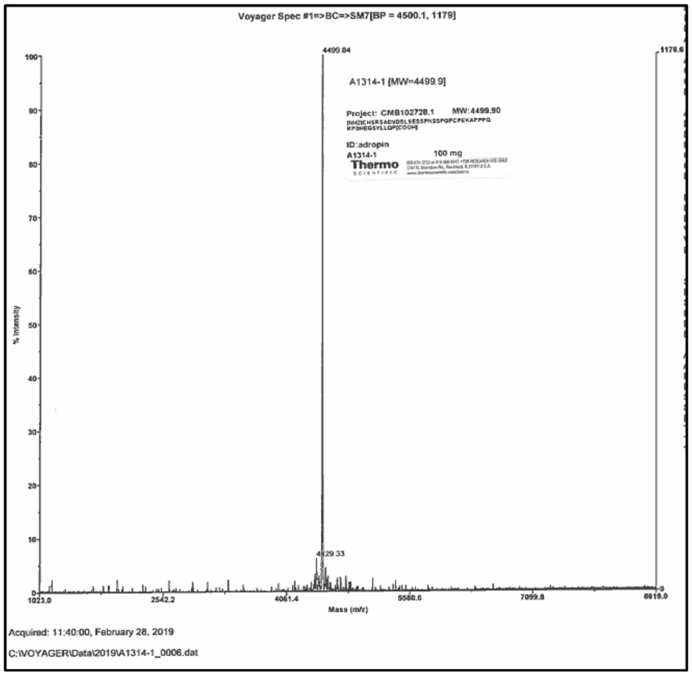


C

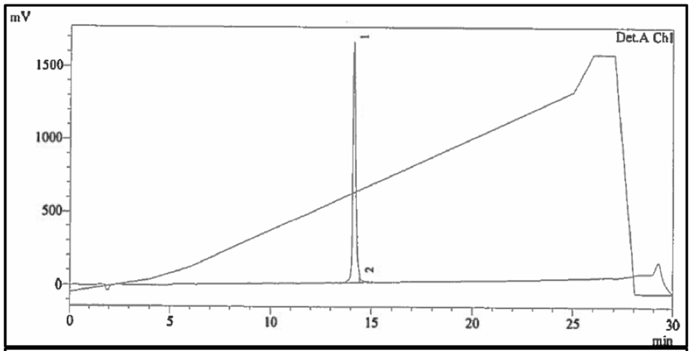


PeakTable

Detector A Ch1 220nm

| Peak# | Ret. Time | Area | Height | Height % | Area % |
|---|---|---|---|---|---|
| 1 | 14.153 | 16181525 | 1651139 | 97.232 | 98.138 |
| 2 | 14.433 | 307072 | 47002 | 2.768 | 1.862 |
| Total | | | | 100.000 | 100.000 |

D

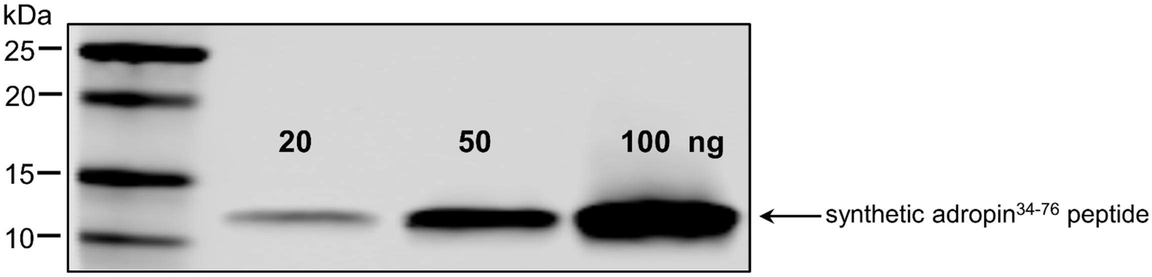


**Supplemental Figure 2. Profile of synthetic adropin$^{34\text{-}76}$ peptide. A,** Amino acid sequence of the synthetic adropin$^{34\text{-}76}$ peptide. **B, C**, HPLC-MS/MS confirmed the purity and molecular identity of the synthetic adropin$^{34\text{-}76}$ peptide. **D**, Western blot shows that the molecular weight for the synthetic adropin$^{34\text{-}76}$ peptide is around 14 kDa.

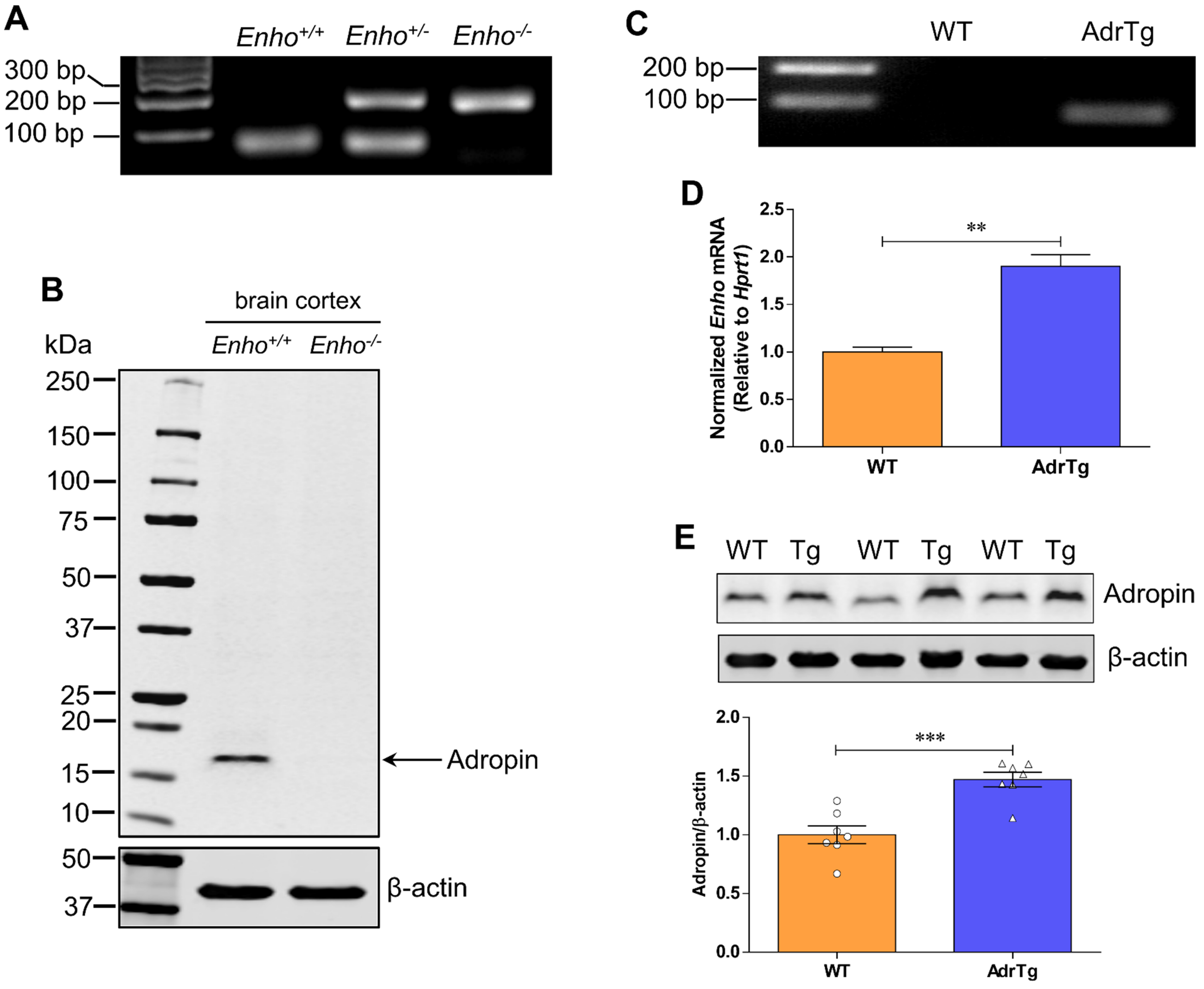


**Supplemental Figure 3. Genotyping and expression analysis of adropin knockout and transgenic mice. A**, An agarose gel electrophoresis shows PCR products for wild-type (*Enho*$^{+/+}$; 72 bp), heterozygous (*Enho*$^{+/-}$; 72 and 170 bp), and homozygous (*Enho*$^{+/-}$; 170 bp) genotypes performed on tail tissue genomic DNA using specific *Enho* wild-type and excised primers. **B,** Adropin monoclonal antibody is highly specific. Full western blot clearly shows a single band appearing at the right molecular weight of adropin (17 kDa) but not in homogenates from adropin knockout (*Enho*$^{-/-}$) mice. **C,** Genotyping of adropin transgenic mice performed on tail tissue genomic DNA by PCR using primers from the human β-actin promoter and *Enho* gene coding region, where the appearance of the 80 bp amplicon confirms the presence of the transgene. **D, E,** Quantitative RT-PCR, and western blot confirm that *Enho* mRNA expression (**D**) and protein levels (**E**) in AdrTg mice are significantly upregulated in the brain compared to the corresponding wild-type littermates. Unpaired t-test, $^{**}P<0.01$, $^{***}P<0.001$. n=3 per group for qRT-PCR, and n=7 per group for western blot.

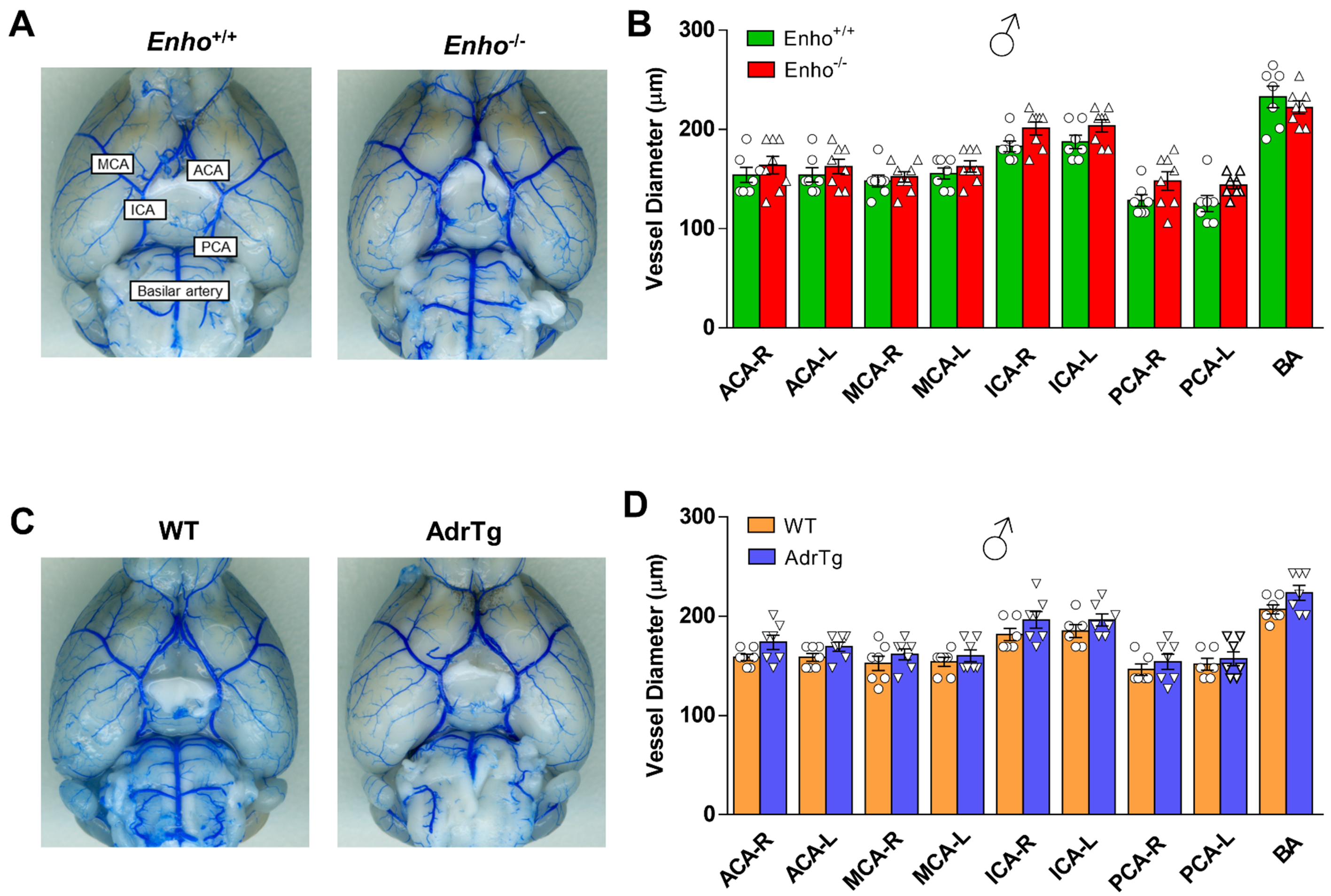


**Supplemental Figure 4. Visualization of the cerebrovascular anatomy in adropin knockout, transgenic mice and their corresponding wild-type littermates.** Intravascular perfusion with latex blue shows permanent staining of large vessels on the ventral surface of brains in adult (10-12 weeks) male *Enho*$^{+/+}$, AdrTg, and their corresponding wild-type littermates. The diameters of the anterior cerebral artery (ACA), middle cerebral artery (MCA) and internal carotid artery (ICA), posterior cerebral artery (PCA), and basilar artery were measured in *Enho*$^{+/+}$, *Enho*$^{-/-}$, WT (the counterpart to AdrTg), and AdrTg mice, and no significant differences were observed between genotypes (**A-D**). Unpaired t-test. *Enho*$^{+/+}$ (n=7), *Enho*$^{-/-}$ (n=8), WT (n=7), AdrTg (n=7).

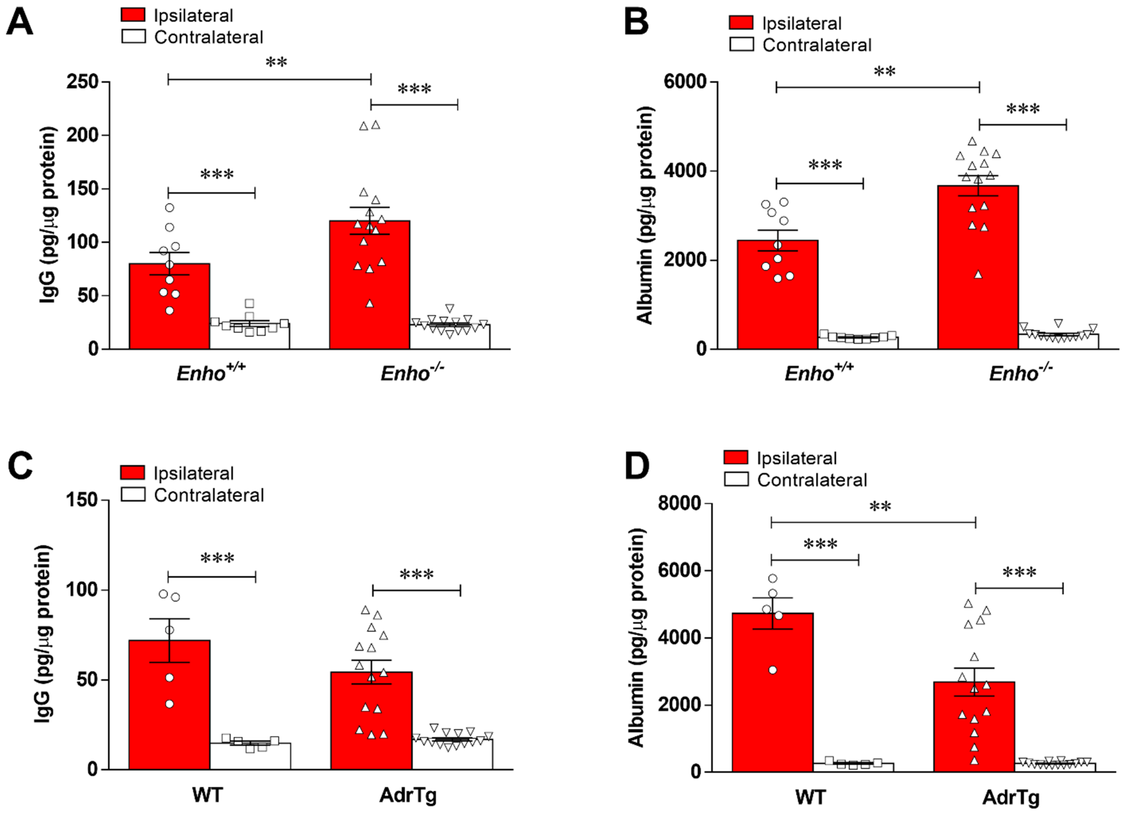


**Supplemental Figure 5. Endogenous adropin is beneficial in the preservation of BBB integrity in male mice subjected to ischemic stroke.** BBB permeability was assessed by measurement of immunoglobulin G (IgG) and albumin at 48h after pMCAO in the ipsilateral and contralateral cerebral cortices. Deficiency of the *Enho* gene (*Enho*$^{-/-}$) resulted in significant extravasation of IgG (**A**) and albumin (**B**) into the ipsilateral cortex compared to *Enho*$^{+/+}$ mice, while overexpressing adropin (AdrTg) reduced the leak of IgG and albumin into ischemic brain compared to the corresponding wild-type mice (**C, D**). Two-way ANOVA with Bonferroni post-tests, ***P*<0.01, ****P*<0.001. *Enho*$^{+/+}$ (n=9), *Enho*$^{-/-}$ (n=14), WT (n=5), AdrTg (n=14).

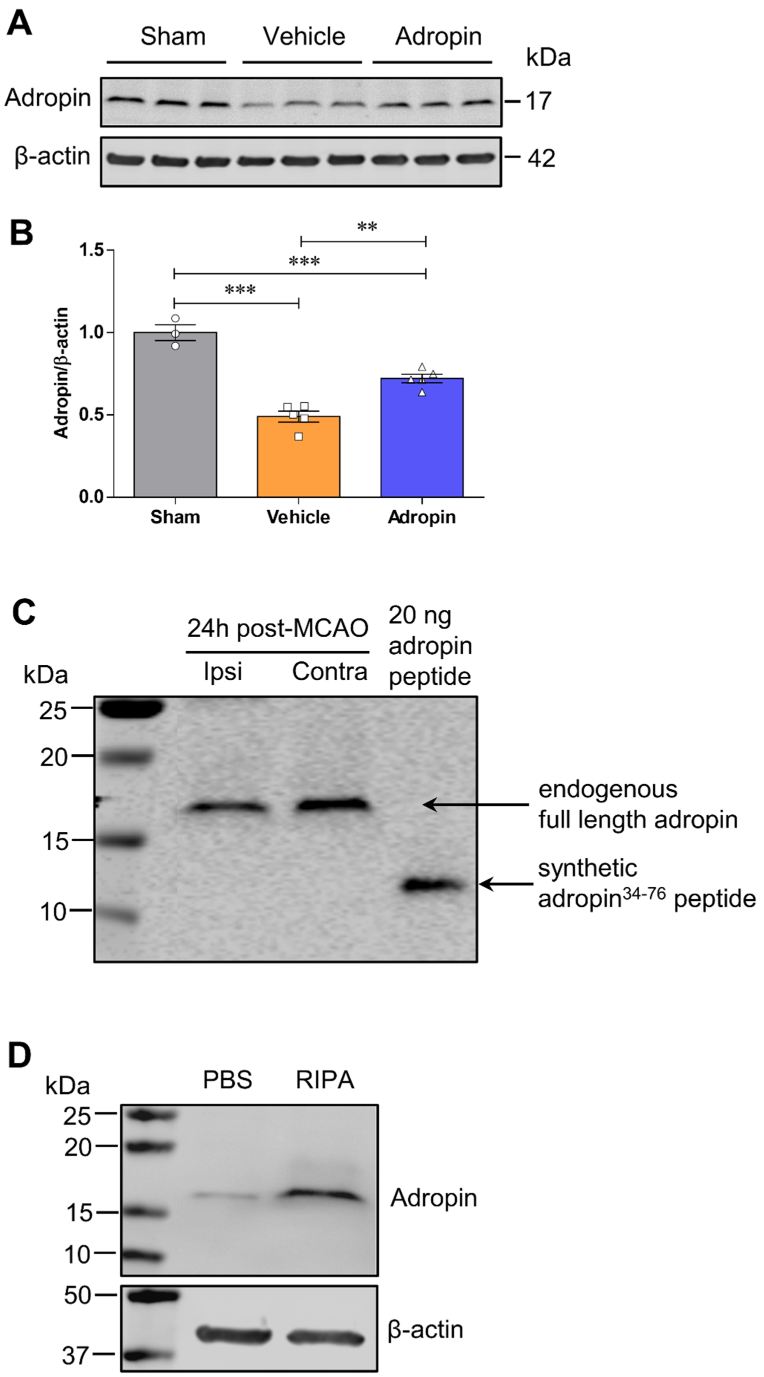


**Supplemental Figure 6. Loss of endogenous adropin in the ischemic brain is attenuated by the treatment of synthetic adropin$^{34\text{-}76}$ peptide. A**, **B**, Adult (10-12 weeks) male mice were given one dose of either vehicle or synthetic adropin$^{34\text{-}76}$ peptide (900 nmol/kg; i.v.) at the onset of cerebral ischemia and euthanized at 24h post-stroke. Endogenous adropin level in homogenates prepared from the ipsilateral cortex was quantified by immunoblotting. Representative Western blots and densitometric analysis showed that adropin treatment significantly attenuated the stroke-induced loss of endogenous adropin levels in the ischemic cerebral cortex. One-way ANOVA with Bonferroni post-tests, **$P$<0.01, ***$P$<0.001. Sham (n=3), Vehicle (n=5), Adropin (n=5). **C**, Representative Western blot for full-length endogenous adropin in the ipsilateral (ischemic) and contralateral (non-ischemic) cerebral cortex at 24h after pMCAO. Synthetic adropin$^{34\text{-}76}$ peptide (20 ng in loading buffer) was run next to the brain samples to compare their molecular weight and running pattern. While the band for synthetic adropin peptide runs at about 14 kDa, the endogenous full-length adropin appears at a higher molecular weight (~17 kDa). **D**, Cerebral cortex from naïve mouse brain was homogenized in either phosphate-buffered saline (PBS) or radioimmunoprecipitation (RIPA) lysis buffer. An equal amount of total protein (50 µg) was separated in a 4-20% SDS-polyacrylamide gel under reducing and denaturing conditions. A dramatic increase in adropin signal was found when proteins were extracted using RIPA buffer containing detergents, suggesting that endogenous adropin in the brain is likely a membrane-bound protein.

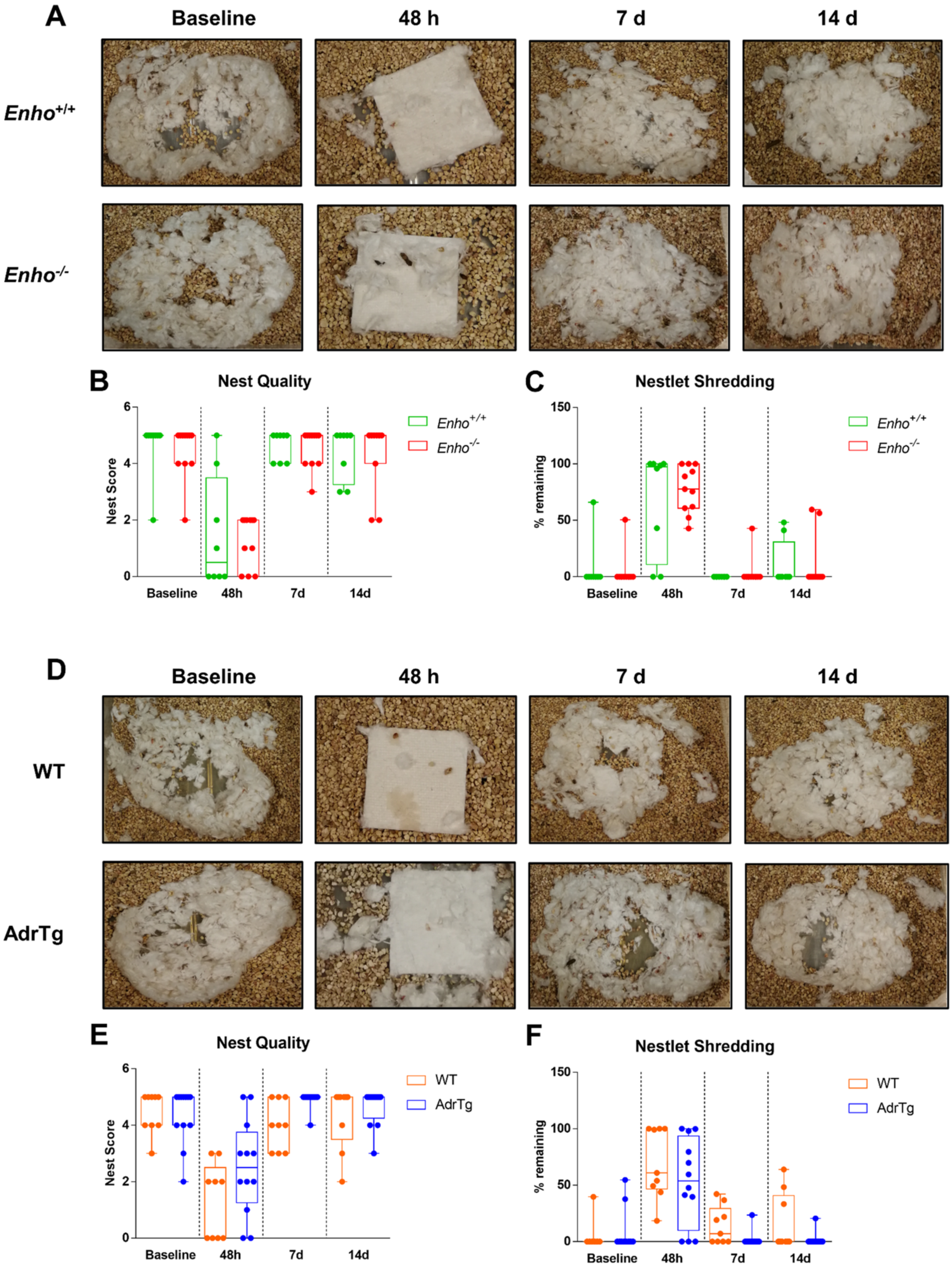


**Supplemental Figure 7. Nesting behavior in mice subjected to pMCAO. A, D**, Representative photographs of nests at baseline and periods of 48 h, 7d and 14d after pMCAO in *Enho*$^{+/+}$, *Enho*$^{-/-}$, WT (the counterpart to AdrTg) and AdrTg mice. **B, C, E, F**, Graphical representations show that stroke-induced dramatic decreases in nesting scores and nestlet shredding ability in all mice at 48h post-MCAO, and the impaired nest-building ability was slightly recovered after 7 days following pMCAO. However, no significant difference was observed in nesting behavior between *Enho*$^{+/+}$ and *Enho*$^{-/-}$ and WT and AdrTg mice. Mann–Whitney Wilcoxon analysis was used for the statistical analysis. *Enho*$^{+/+}$ (n=8), *Enho*$^{-/-}$ (n=11), WT (n=9), AdrTg (n=12).

**Supplemental Table 1. Physiological parameters and blood pressure of adropin transgenic, knockouts and their respective wild-type control mice**

| | **WT (n=9)** | **AdrTg (n=9)** | ***Enho*$^{+/+}$ (n=6)** | ***Enho*$^{-/-}$ (n=7)** |
|---|---|---|---|---|
| pH | 7.35±0.09 | 7.32±0.07 | 7.38±0.07 | 7.31±0.08 |
| $PCO_2$ (mmHg) | 34.17±5.65 | 38.40±4.75 | 33.92±8.30 | 37.93±4.34 |
| $PO_2$ (mmHg) | 299.89±66.77 | 260.27±134.84 | 320.40±58.90 | 358.71±103.86 |
| $HCO_3^-$ (mmol/L) | 18.90±2.17 | 20.05±2.01 | 19.82±2.42 | 19.29±3.06 |
| $Na^+$ (mmol/L) | 144.44±1.81 | 142.27±2.85 | 144.00±5.05 | 144.43±0.53 |
| $K^+$ (mmol/L) | 4.27±0.42 | 4.44±0.21 | 4.50±0.67 | 4.20±0.27 |
| $Ca^{2+}$ (mmol/L) | 1.15±0.10 | 1.14±0.13 | 1.15±0.14 | 1.18±0.06 |
| Glucose (mg/dL) | 215.09±36.94 | 227.93±26.85 | 231.96±40.21 | 230.46±49.30 |
| Hematocrit (%) | 37.33±1.00 | 36.82±1.73 | 33.60±3.44 | 37.00±1.83 |
| Hemoglobin (g/dL) | 12.69±0.36 | 12.23±0.59 | 11.42±1.18 | 12.59±0.61 |
| SBP (mmHg) | 111.4±15.9 | 112.2±12.7 | 113.8±5.3 | 116.6±4.9 |
| DBP (mmHg) | 73.6±12.1 | 76.4±9.7 | 79.0±6.4 | 82.7±12.5 |
| MAP (mmHg) | 86.1±12.8 | 88.4±10.1 | 90.3±3.6 | 93.6±9.8 |
| HR (beats/min) | 397±62 | 357±68 | 345±20 | 367±97 |

SBP: systolic blood pressure, DBP: diastolic blood pressure, MAP: mean arterial pressure, HR: heart rate. WT (wild-type) was the control group for the adropin transgenic (AdrTg) mouse line. Data represented as mean ± SD. Unpaired Student's *t* test was performed between WT and AdrTg, *Enho*$^{+/+}$ and *Enho*$^{-/-}$ mice.

**Supplemental Table 2. Antibodies used for western blot and immunohistochemistry**

| Antibodies | Source | Identifier | Dilution |
|---|---|---|---|
| mouse anti-Adropin | Cayman Chemical | CAT#: 14117 | 1:1000 |
| rabbit anti-phospho-eNOS | Cell Signaling Technology | CAT#: 9571 | 1:500 |
| rabbit anti-phospho-Akt | Cell Signaling Technology | CAT#: 4060 | 1:1000 |
| rabbit anti-phospho-$ERK_{1/2}$ | Cell Signaling Technology | CAT#: 4370 | 1:1000 |
| rabbit anti-eNOS | Santa Cruz Biotechnology | CAT#: sc-654 | 1:500 |
| rabbit anti-Akt | Cell Signaling Technology | CAT#: 4691 | 1:1000 |
| mouse anti-$ERK_{1/2}$ | Cell Signaling Technology | CAT#: 4696 | 1:1000 |
| goat anti-CD31/PECAM-1 | Novus Biologicals | CAT#: 77699 | 1:2000 |
| rabbit anti-ZO-1 | Invitrogen | CAT#: 61-7300 | 1:500 |
| rabbit anti-Occludin | Abcam | CAT#: ab167161 | 1:1000 |
| rabbit anti-Claudin-5 | Abcam | CAT#: ab124284 | 1:2000 |
| rabbit anti-MMP-9 | Santa Cruz Biotechnology | CAT#: sc-6841-R | 1:500 |
| mouse anti-gp91phox | BD Biosciences | CAT#: 611414 | 1:500 |
| mouse anti-4-HNE | Abcam | CAT#: ab48506 | 1:200 |
| mouse anti-Ly-6B.2 | Bio-Rad | CAT#: MCA771GA | 1:500 |
| mouse anti-β-actin | Sigma-Aldrich | CAT#: A1978 | 1:10000 |
| IRDye 800CW Goat anti-rabbit IgG | LI-COR Biosciences | CAT#: 926-32211 | 1:30000 |
| IRDye 800CW Goat anti-mouse IgG | LI-COR Biosciences | CAT#: 926-32210 | 1:30000 |
| IRDye 680LT Donkey anti-mouse IgG | LI-COR Biosciences | CAT#: 926-68022 | 1:40000 |
| IRDye 800CW Donkey anti-goat IgG | LI-COR Biosciences | CAT#: 926-32214 | 1:30000 |
| rat anti-CD31/PECAM-1 | Novus Biologicals | CAT#: NB600-1475 | 1:100 |
| rabbit anti-NeuN | Novus Biologicals | CAT#: NBP1-77686 | 1:100 |
| rabbit anti-GFAP | Dako | CAT#: Z0334 | 1:100 |
| rabbit anti-Iba1 | Wako | CAT#: 019-19741 | 1:100 |
| Goat anti-mouse IgG Alexa Fluor 594 | ThermoFisher Scientific | CAT#: A11032 | 1:250 |
| Goat anti-rat IgG Alexa Fluor 488 | ThermoFisher Scientific | CAT#: A11006 | 1:250 |
| Goat anti-rabbit IgG Alexa Fluor 488 | Jackson ImmunoResearch | CAT#: 111-545-144 | 1:250 |

**Supplemental Table 3. Primers used for genotyping and qRT-PCR**

| Purpose | Allele/Gene | Type | Primer sequence (5'-3') | Product size (bp) |
|---|---|---|---|---|
| Genotyping | wild-type | Forward | GGAAGATGTACGTTCAGACACACA | 72 |
| | | Reverse | GTGTGCCTAGGTGTACCTTTCC | |
| | excised | Forward | ACGTTCAGACACACACCTGATC | 170 |
| | | Reverse | CTACAATCTAGGCTGCTGGGT | |
| | mutant | Forward | GGCTATTCTCGCAGGATCAGT | 80 |
| | | Reverse | GAGAGCCCCTTGGGAGATG | |
| qRT-PCR | *Enho* | Forward | ATGGCCTCGTAGGCTTCTTG | N/A |
| | | Reverse | GGCAGGCCCAGCAGAGA | |
| | *Hprt1* | Forward | CCCCAAAATGGTTAAGGTTGC | N/A |
| | | Reverse | AACAAAGTCTGGCCTGTATCC | |

#### Full unedited gels for Figure 1A

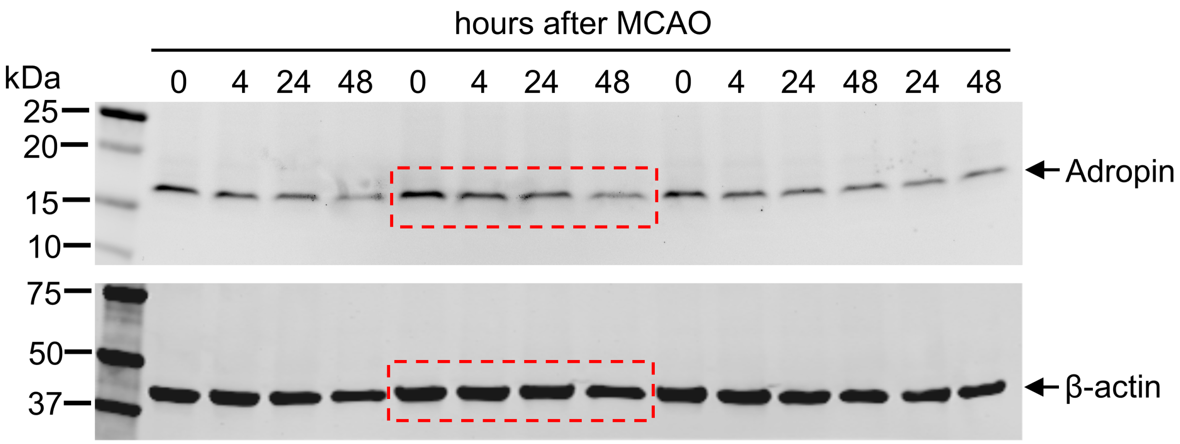

#### Full unedited gels for Figure 4D

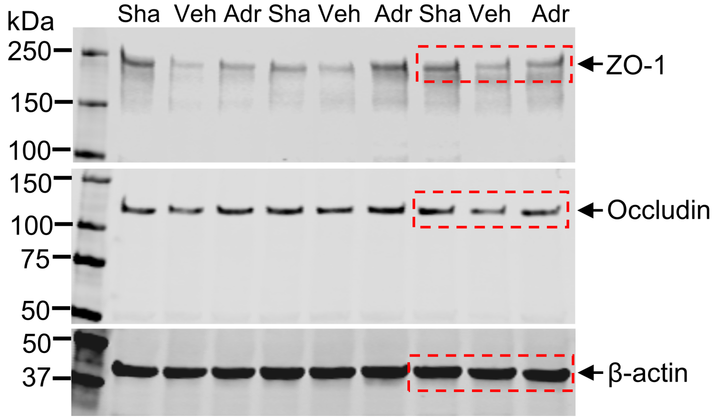


#### Full unedited gels for Figure 4G

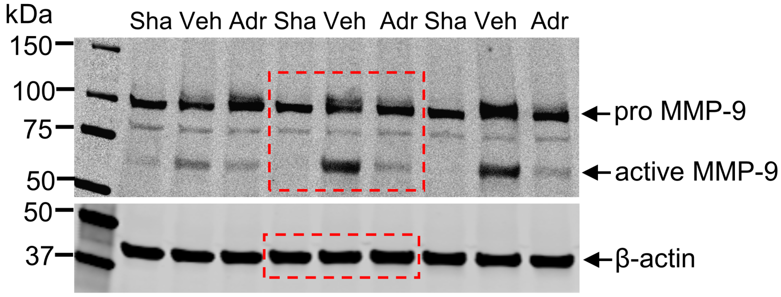


#### Full unedited gels for Figure 4J

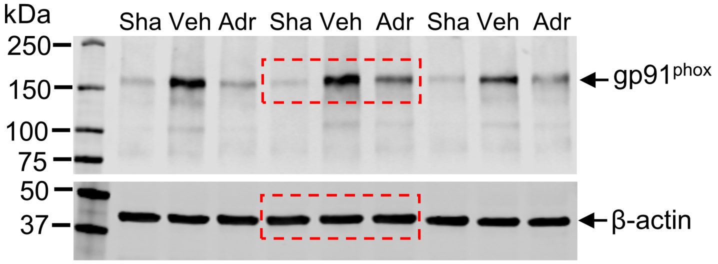


#### Full unedited gels for Figure 4L

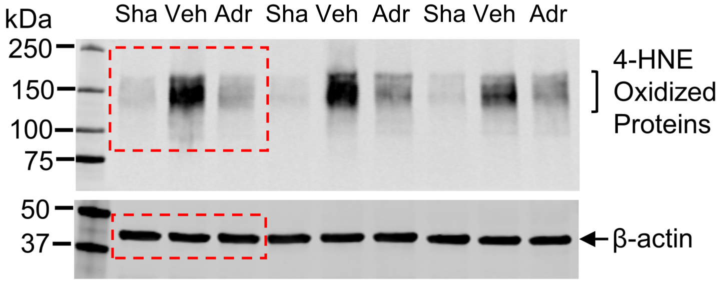


#### Full unedited gels for Figure 4N

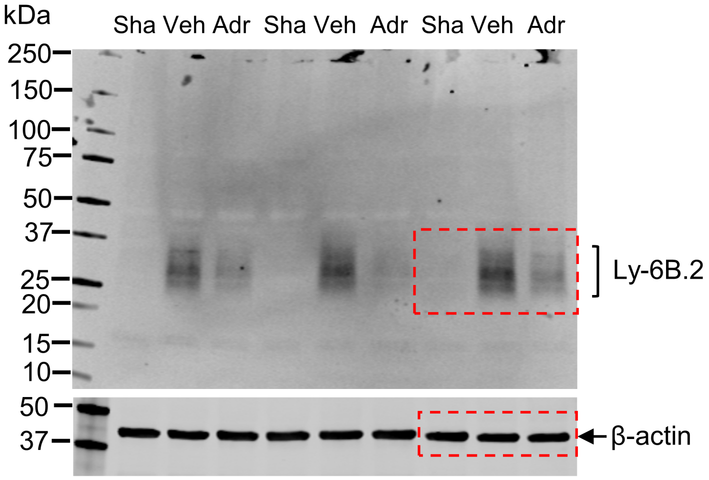

#### Full unedited gels for Figure 5A

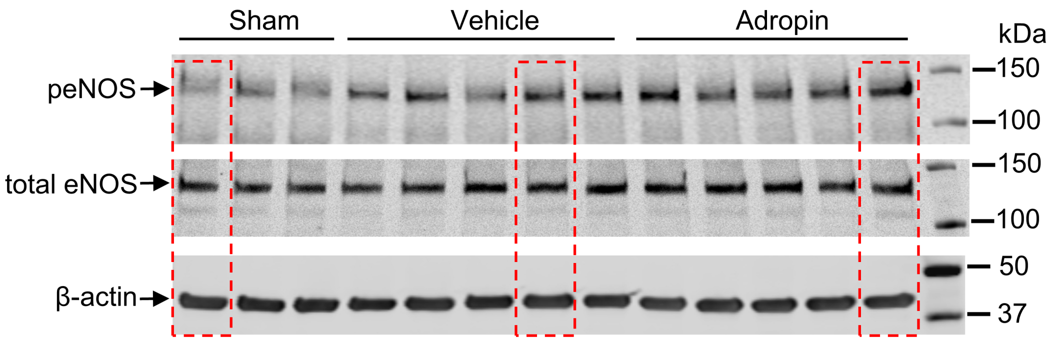

#### Full unedited gels for Figure 6D

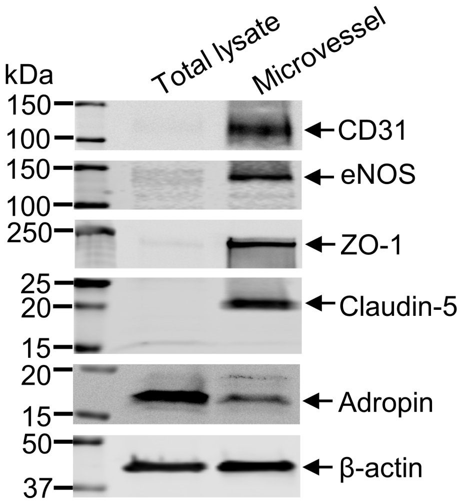


#### Full unedited gels for Figure 6E

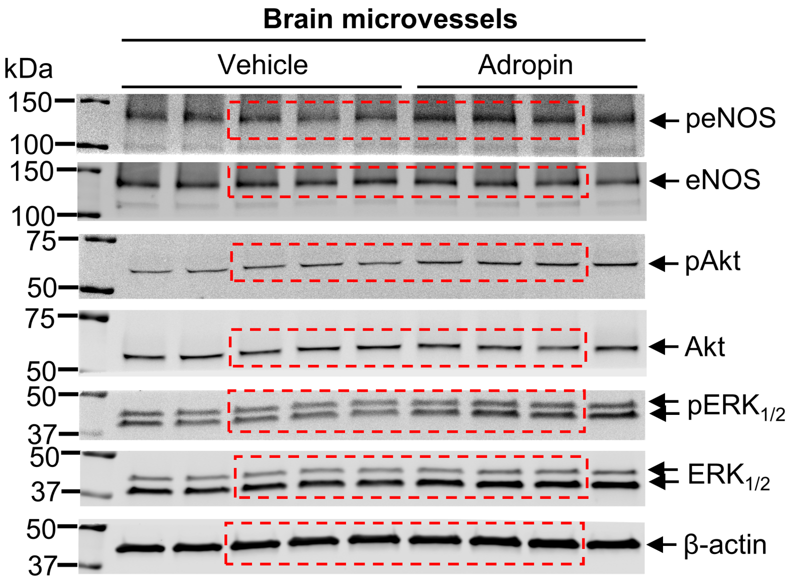


#### Full unedited gels for Figure 6J

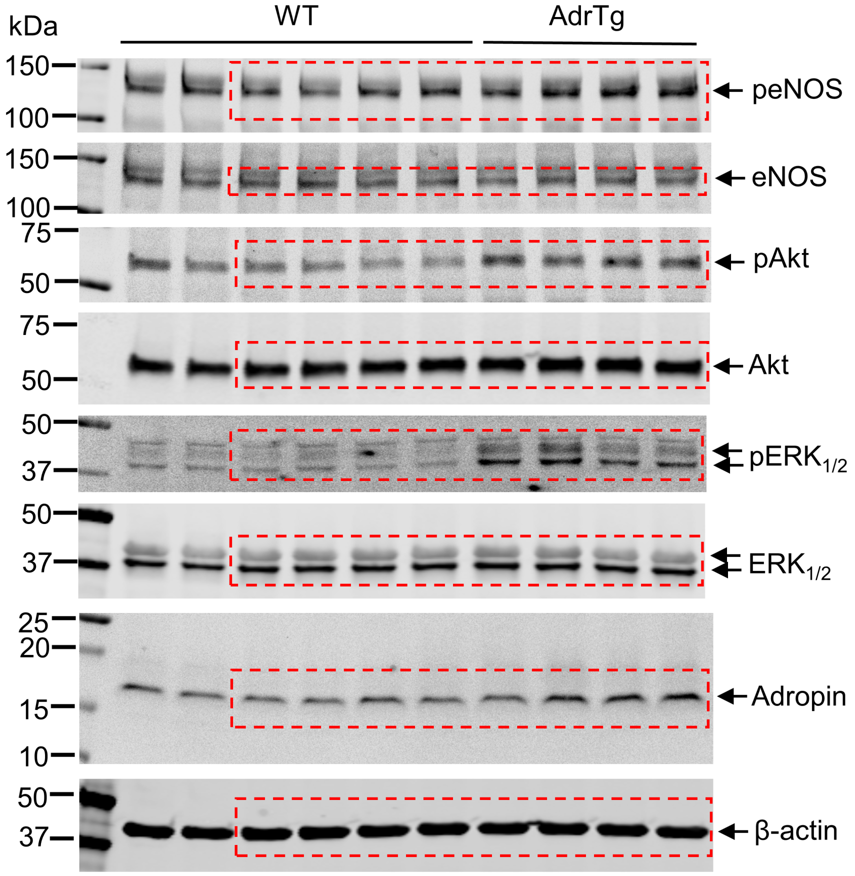

#### Full unedited gels for Supplemental Figure 1C

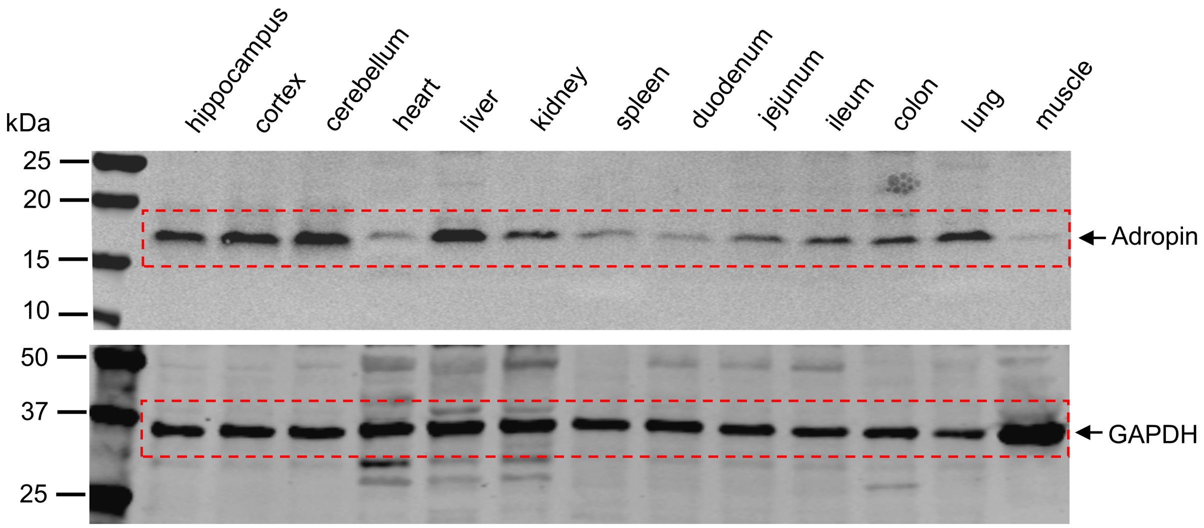

#### Full unedited gel for Supplemental Figure 2D

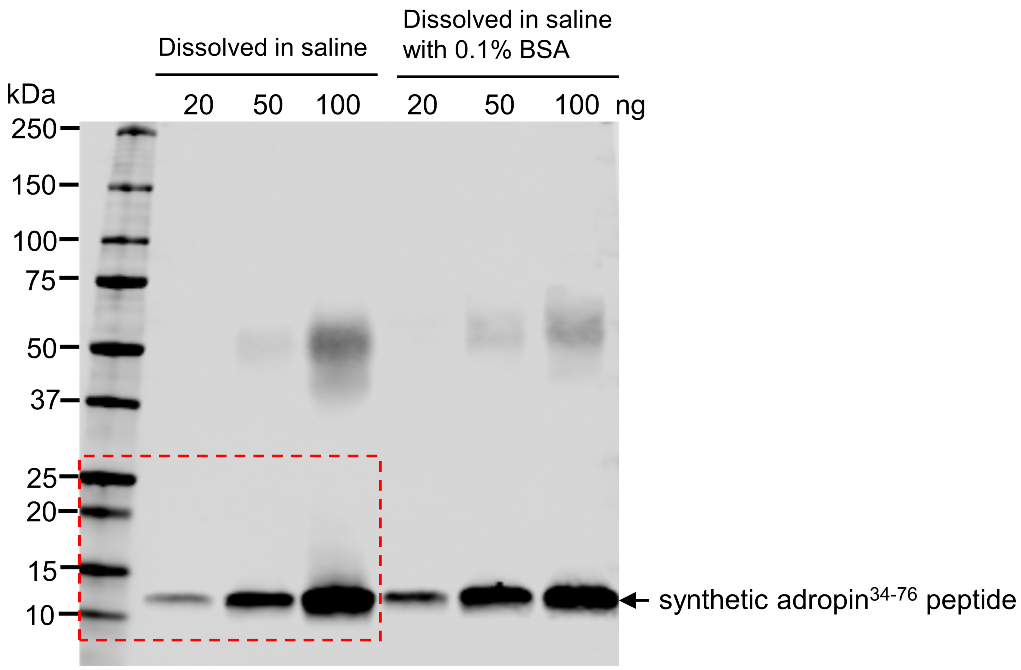

### Full unedited gel for Supplemental Figure 3E

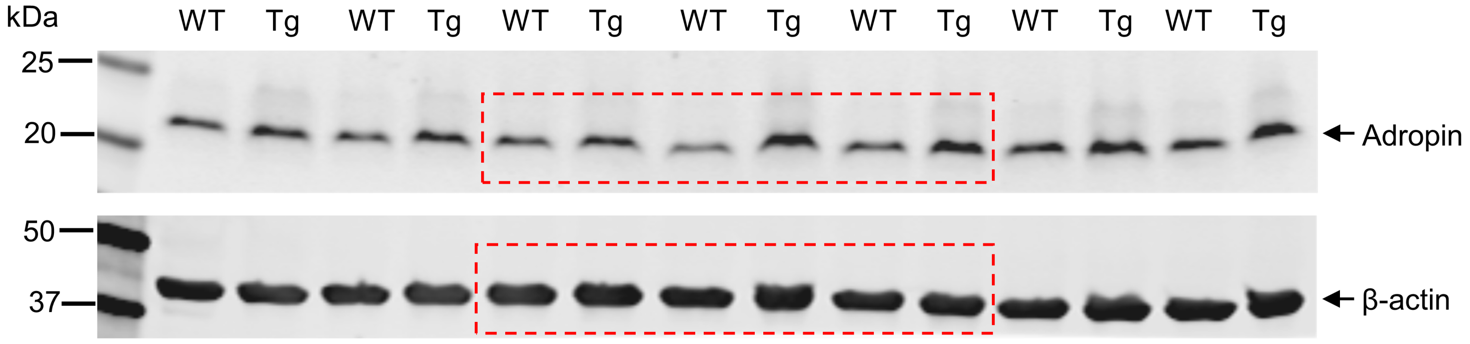

#### Full unedited gels for Supplemental Figure 6A

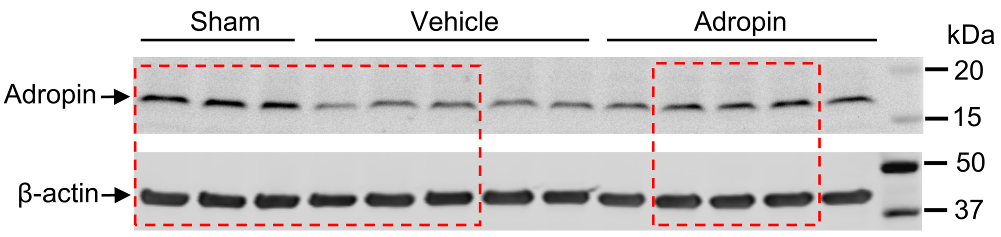


#### Full unedited gel for Supplemental Figure 6C

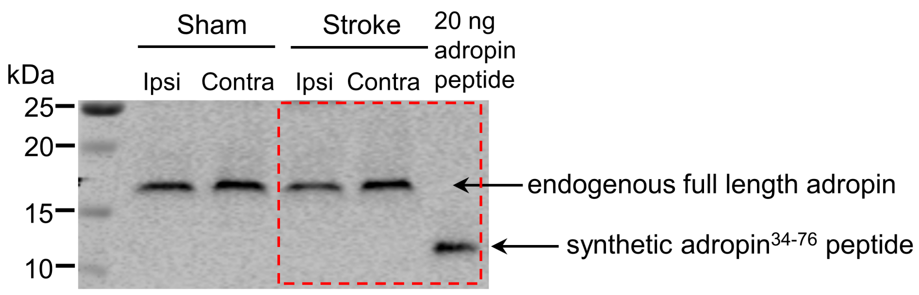


#### Full unedited gel for Supplemental Figure 6D

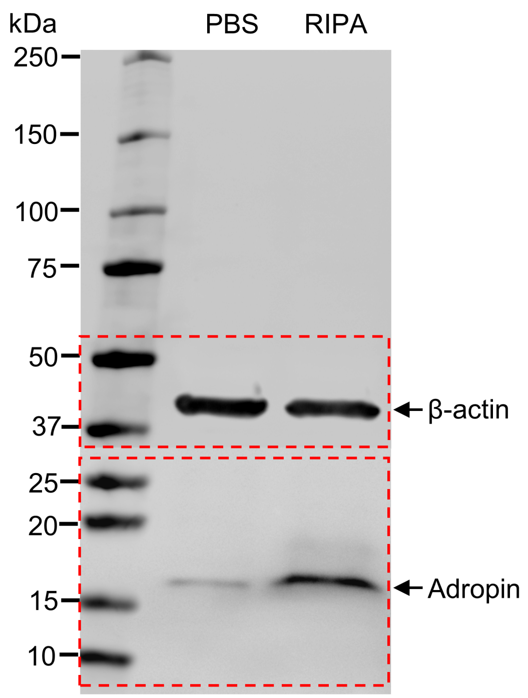
